# DRP1 IS REQUIRED FOR AGRP NEURONAL ACTIVITY AND FEEDING

**DOI:** 10.1101/2020.10.11.335224

**Authors:** Sungho Jin, Nal Ae Yoon, Zhong-Wu Liu, Tamas L. Horvath, Jung Dae Kim, Sabrina Diano

## Abstract

The hypothalamic orexigenic Agouti-related peptide (AgRP)-expressing neurons are crucial for the regulation of whole-body energy homeostasis. Here, we show that fasting-induced AgRP neuronal activation is associated with dynamin-related peptide 1 (Drp1)-mediated mitochondrial fission and mitochondrial fatty acid utilization in AgRP neurons. In line with this, mice lacking Drp1 in adult AgRP neurons (*Drp1*^*AgRPKO*^) show decrease in fasting-or ghrelin-induced AgRP neuronal activity and feeding and exhibited a significant decrease in body weight, fat mass, and feeding accompanied by a significant increase in energy expenditure. In support of the role for mitochondrial fission and fatty acids oxidation, *Drp1*^*AgRPKO*^ mice showed attenuated palmitic acid-induced mitochondrial respiration. Altogether, our data revealed that mitochondrial dynamics and fatty acids oxidation in hypothalamic AgRP neurons is a critical mechanism for AgRP neuronal function and associated behavior.

## Introduction

The central nervous system (CNS) regulates whole-body energy metabolism through multiple neuronal networks (Myers and Olson, 2012, Diano, 2013). The hypothalamus has been considered a key area of the brain in regulating metabolism via the ability of hypothalamic neurons to sense, integrate, and respond to fluctuating metabolic signals (Sandoval et al., 2009, Coll and Yeo, 2013). The hypothalamic arcuate nucleus (ARC) contains two distinct neuronal subpopulations that produce either orexigenic neuropeptides agouti-related peptide (AgRP) and neuropeptide-Y (NPY), or anorexigenic neuropeptides including alpha-melanocyte stimulating hormone (α-MSH) derived from proopiomelanocortin (POMC) (Ollmann et al., 1997, Batterham et al., 2002, Roh et al., 2016). The anatomical location of the hypothalamic ARC allows these neurons to rapidly respond to fluctuations of numerous circulating metabolic signals, including nutrients and hormones (Gao and Horvath, 2007). However, the intracellular mechanisms underlying their ability to sense circulating signals, and, specifically nutrients, remain to be elucidated.

Mitochondria are the main powerhouse of the cell by producing adenosine triphosphate (ATP) (Mattson et al., 2008)(Picard et al., 2016). Neurons rely on mitochondrial electron transport chain and oxidative phosphorylation to meet their high energy demands (Belanger et al., 2011). In addition, mitochondria are highly dynamic organelles able to change their morphology and location according to the needs of the cell (Chan, 2006). The ability of mitochondria to change their morphological characteristics in response to the metabolic state to match with the needs of the cells occurs through fusion and fission events, process defined as mitochondrial dynamics. Mitochondrial morphological changes are associated with several proteins, including mitofusin 1 and 2 (MFN1 and MFN2) in the mitochondrial outer membrane and optic atrophy-1 (OPA1) in the mitochondrial inner membrane for mitochondrial fusion (Youle and van der Bliek, 2012, Kasahara and Scorrano, 2014), whereas mitochondrial fission is regulated by the activity of the dynamin-related protein 1 (Drp1), which is recruited to the mitochondrial outer membrane to interact with mitochondrial fission factor (Mff) and mitochondrial fission 1 (Fis1) (Loson et al., 2013).

Previous studies from our laboratory has shown that NPY/AgRP neuronal activation is associated with changes in mitochondrial morphology and density during fasting or after ghrelin administration (Coppola et al., 2007; Andrews et al., 2008; Dietrich et al., 2013), suggesting that changes in mitochondrial dynamics may play a role in the regulation of neuronal activation of these neurons (Nasrallah and Horvath, 2014). In addition, in diet-induced obesity (DIO) mouse model, silent NPY/AgRP neurons showed mitochondrial dynamics leaning towards mitochondrial fusion (Dietrich et al., 2013). In the present study we interrogated the relevance of mitochondrial fission in AgRP neurons in relation to fuel availability.

## Results

### Fasting induces mitochondrial fission in AgRP neurons

Recent studies have demonstrated that hypothalamic mitochondrial dynamics play a critical role in regulating nutrient sensing (Dietrich et al., 2013; Schneeberger et al., 2013; Toda et al., 2016; Santoro et al., 2017). Using electron microscopy, we observed that compared to feeding (0.174 ± 0.007 µm^2^, p<0.0001; Figure 1a,c), fasting resulted in a significant decrease in mitochondrial size (0.130 ± 0.005 µm^2^, Figure 1b,c) in AgRP neurons together with a significant increase in mitochondrial density (0.551 ± 0.032 mitochondria/ µm^2^ of cytosol in fasting vs 0.423 ± 0.026 mitochondria/ µm^2^ of cytosol in feeding; ; p=0.0031; Figure 1d). This was associated wih a decrease in mitochondrial aspect ratio (1.629 ± 0.020 in fasting vs 1.769 ± 0.049 in feeding; p=0.0064; Figure 1e). However, total mitochondrial coverage in the cytosol (Figure 1f) in AgRP neurons was not altered between fed (7.237 ± 0.461 % of cytosol) and fasted mice (6.830 ± 0.363 % of cytosol; p=0.4853). These observations indicate that food deprivation promotes mitochondrial fission in AgRP neurons.

**Figure 1.**
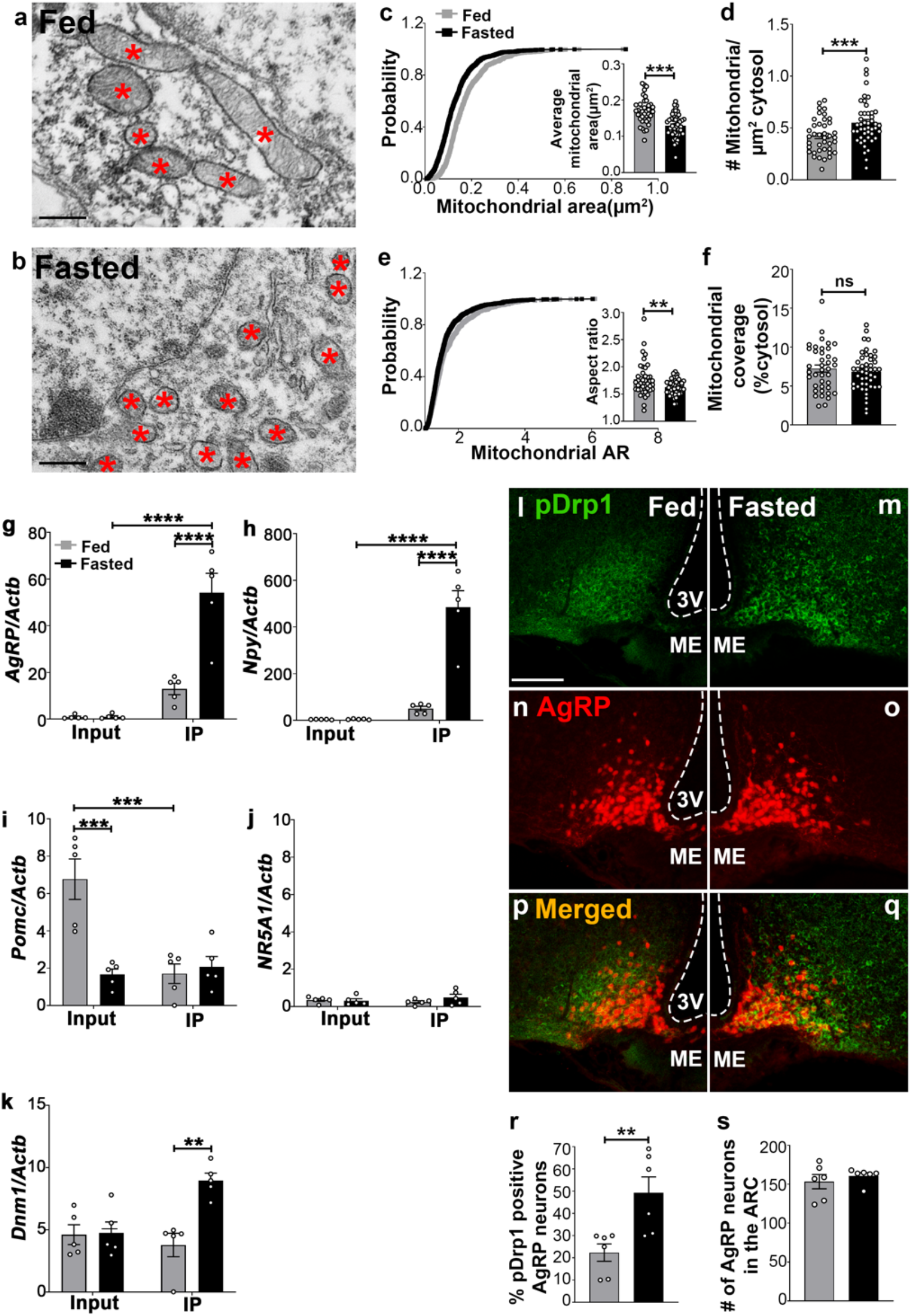
Fasting induces mitochondrial fission and activation of Drp1 in AgRP neurons. **(a-b)** Representative electron micrographs showing mitochondria (asterisks) in an AgRP neuron of 5 months old fed (a) and the fasted male mouse (b). Scale bar represents 500 nm. **(c-f)** Cumulative probability distribution of cross-sectional mitochondria area and average mitochondrial area (c), mitochondrial density (d), a cumulative probability distribution of mitochondrial aspect ratio (e) and mitochondrial coverage (f) in AgRP neurons from fed and fasted male mice (fed mice, n=779 mitochondria/39 AgRP neurons/4 mice; fasted mice, n=1559 mitochondria/47 AgRP neurons/6 mice). Data are presented as mean ± SEM. **P<0.01; ****P<0.0001 by two-tailed Student’s *t*-test. ns=not significant. **(g-k**) Real-time PCR data showing relative mRNA levels of *AgRP* (g), *Npy* (h), *Pomc* (i), *NR5A1* (*Sf1*, j), and *Dnm1* (*Drp1*, k) in total lysate of hypothalami (Input) and isolated RNA bound to the ribosomes of the hypothalamic AgRP neurons (IP) from 3 months old fed or fasted mice (n=5/ group). Three animals were pooled for each n. Data are presented as mean ± SEM. **P<0.01; ***P<0.001; ****P<0.0001 by two-way ANOVA with Tukey’s post hoc analysis for multiple comparisons. **(l-q)** Representative micrographs showing immunostaining for phosphorylated Drp1 (at serine 616; pDrp1; green, l-m) and *td*Tomato (red, representing AgRP, n-o) and merged (p-q) in the hypothalamic ARC of 5 months old fed and fasted male mice. Scale bar represents 100 µm. 3v=third ventricle; ARC=arcuate nucleus; ME=median eminence. **(r)** Graph showing the percentage of AgRP neurons immunopositive for pDrp1 (n=6 mice/ group). Data are presented as mean ± SEM. **P<0.01 by two-tailed Student’s *t*-test. **(s)** Graph showing no difference in total AgRP cell number between fed and fasted male mice (n=6 mice/ group). Data are presented as mean ± SEM. P=0.4711 by two-tailed Student’s *t*-test.

### Cell-type-specific transcriptomic analysis of AgRP neurons in the hypothalamus

We next performed transcriptomic profiling using ribosomal tagging strategy to analyze AgRP neuron-specific mRNA expression levels in fed and fasted *AgRP-cre:ER*^*T2*^; *RiboTag* mice. Compared to the input RNA, a 55 to 107 fold enrichment of transcripts for AgRP and NPY expressed in the immunoprecipitated RNA (IP), respectively, was observed in fasted *AgRP-cre:ER*^*T2*^; *RiboTag* mice (*AgRP*= input=0.974 ± 0.441, n=5; IP=54.120 ± 8.300, n=5; p<0.0001, Figure 1g; *Npy=* input=4.516 ± 0.892, n=5; IP=485.424 ± 69.891, n=5; p<0.0001, Figure 1h). Conversely, a significant decrease of transcript for POMC in the IP was found in fed *AgRP-cre:ER*^*T2*^; *RiboTag* mice (1.703 ± 0.523, n=5; p=0.0004, Figure 1i) compared to the input RNA (6.768 ± 1.084, n=5). Moreover, marginal expression of *Nr5a1* was detected in the input RNA (fed=0.350 ± 0.074, n=5; fasted=0.308 ± 0.105, n=5, Figure 1j) and the IP (fed=0.241 ± 0.067, n=5; fasted=0.486 ± 0.173, n=5; p=0.4424, Figure 1j), validating the arcuate AgRP neuronal isolation protocol. In support of our mitochondrial morphology data, quantitative real time-PCR (qRT-PCR) analyses of ribosome-bound mRNAs revealed that the transcription level of *Drp1* (fed=4.754 ± 0.875, n=5; fasted=8.971 ± 0.587, n=5; p=0.0018, Figure 1k) was significantly upregulated in AgRP neurons of fasted mice compared to fed mice.

### Increased Drp1 activation in AgRP neurons of fasted mice

Mitochondria fission is mediated by Drp1, which is recruited to the outer membrane of mitochondria to promote mitochondrial fragmentation in a GTPase-dependent manner followed by its phosphorylation at serine 616 site (Liesa et al., 2009). To examine whether food deprivation is associated with changes in activated Ser616 phosphorylation of Drp1 (pDrp1) levels, we assessed the distribution of pDrp1 immunoreactivity in AgRP neurons in fed and fasted mice. We found that percent of AgRP neurons expressing pDrp1 were significantly increased in fasting (49.2 ± 7.228 % of AgRP neurons, n=6; Figure m,o,q,r) compared to the fed condition (22.33 ± 3.921 % of AgRP neurons, n=6, p=0.0085, Figure 1l,n,p,r). No changes in AgRP cell number were observed between fed (152.2 ± 11.14; n=5; Figure 1s) and fasted mice (160.8 ± 4.028, n=6; p=0.4527, Figure 1s). These data suggest that activation of AgRP neurons in fasting state is closely associated with increased Drp1 activation and, thus, mitochondrial fission, suggesting that Drp1-mediated mitochondrial dynamics may play a role in the regulation of AgRP neuronal activity in fasting state.

### Fasting-induced mitochondrial β-oxidation in the hypothalamic neurons

The hypothalamus is a key region in the control of energy metabolism via the ability of hypothalamic neurons to response to numerous metabolic signals, including nutrients (Jin and Diano, 2018). It has been proposed that hypothalamic availability of free fatty acids controls food intake (Lam et al., 2005) and AgRP function (Andrews et al., 2008). To investigate the effect of fatty acids on mitochondrial β-oxidation in hypothalamic neurons, we assessed palmitic acid (PA)-induced mitochondrial oxygen consumption rate in primary hypothalamic neuronal cell cultures. A significant difference in PA-induced mitochondrial maximal oxygen consumption rate were observed in primary hypothalamic neurons according to the amount of glucose present in the culture (Figure 2). In high glucose concentration, the rate of PA-induced oxygen consumption was significantly lower (Figure 2a-b) compared to that measured in low glucose (Figure 2c-d). Furthermore, under both high and low glucose conditions, a significant decrease in maximal oxygen consumption rate was observed by the addition of the etomoxir, inhibitor of carnitine palmitoyltransferase-1 (CPT1), transporter of fatty acids into the mitochondria. Together, these data suggest that similar to fasting state, when glucose levels are low hypothalamic neurons utilize fatty acids, such as palmitate, as substrates for mitochondrial respiration.

**Figure 2.**
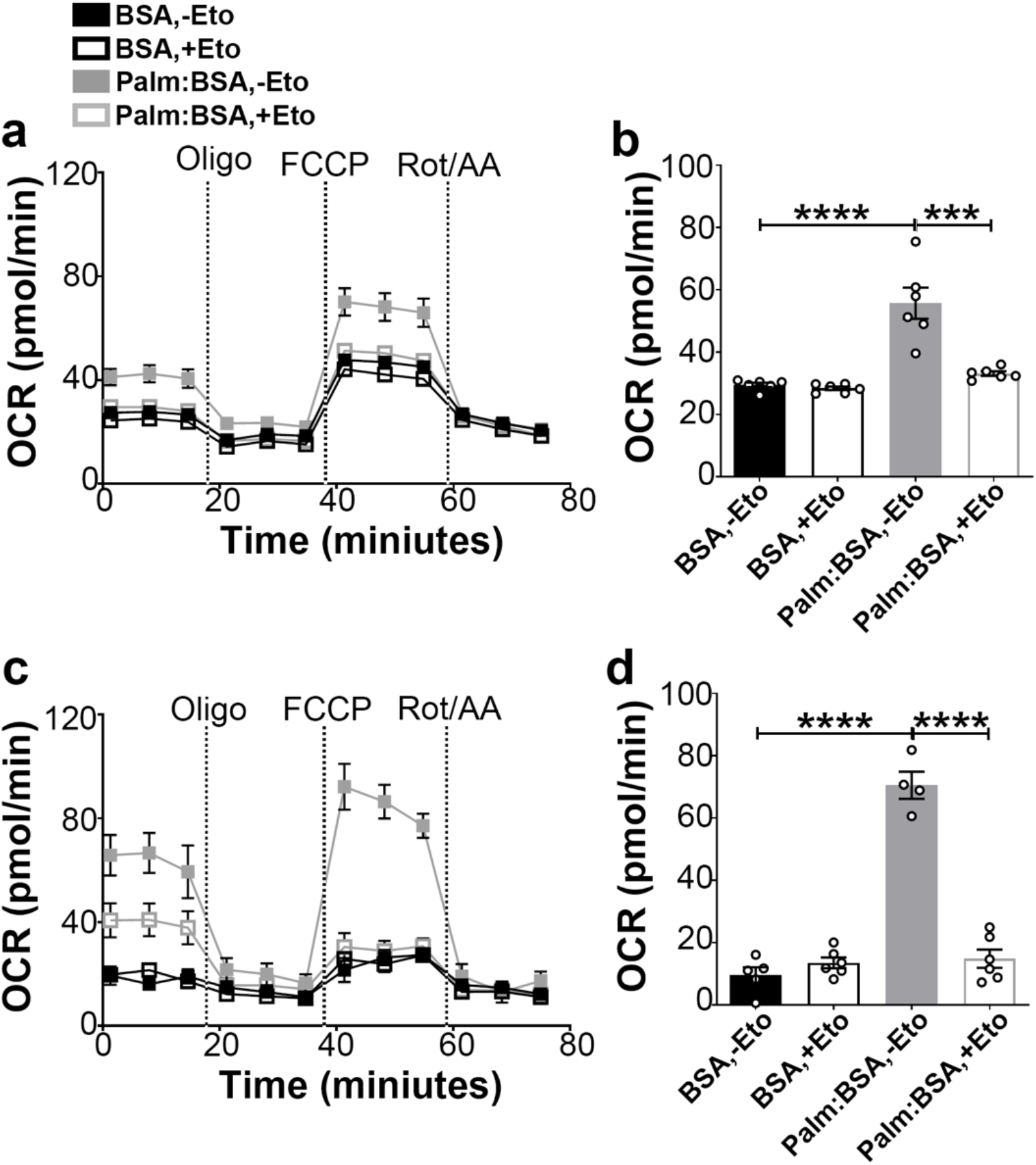
Fasting-induced β-oxidation in the hypothalamic neurons. **(a)** Graphs showing oxygen consumption rate (OCR) under 2.5 mM glucose incubation with or without palmitate-BSA (200 μM) and with or without etomoxir (40 μM) in primary hypothalamic neuronal culture (n=6-8/ group). **(b)** Graph showing the quantification of OCR showed in panel (a) in primary hypothalamic neuronal culture. Data are presented as mean ± SEM. *P<0.05; ***P<0.001; ****P<0.0001 by two-way ANOVA with Tukey’s post hoc analysis for multiple comparisons. **(c)** Graphs showing OCR under low glucose (0.5 mM) with or without palmitate-BSA (200 μM) and with or without etomoxir (40 μM) in primary hypothalamic neuronal culture (n=6-8/ group). **(d)** Graph showing the quantification of OCR shown in panel c in primary hypothalamic neuronal culture. Data are presented as mean ± SEM. ***P<0.001; ****P<0.0001 by two-way ANOVA with Tukey’s post hoc analysis for multiple comparisons.

### Inducible deletion of Drp1 in AgRP neurons

Next, to investigate the physiological functions of Drp1 in adult AgRP neurons, we generated mice with selective and inducible deletion of Drp1 in AgRP neurons (Figure supplement 1a). *AgRP-cre:ER*^*T2*^;*tdTomato* mice (kindly provided by Dr. Joel Elmquist at UTSW) were crossed with Drp1 floxed mice (*Drp1*^*fl/fl*^) (Kageyama et al., 2014). As control groups, *Drp1*^*+/+*^*-AgRP-cre:ER*^*T2*^;*tdTomato* mice were injected with tamoxifen and *Drp1*^*fl/fl*^*-AgRP-cre:ER*^*T2*^;*tdTomato* were mice injected with corn oil. *Drp1*^*fl/fl*^*-AgRP-cre:ER*^*T2*^;*tdTomato* mice (*Drp1*^*AgRPKO*^ mice) were injected with tamoxifen to induce mature-onset deletion of Drp1 in AgRP neurons. To validate our animal model, we analyzed and found limited pDrp1 expression in the AgRP neurons of fasted *Drp1*^*fl/fl*^*-AgRP-cre:ER*^*T2*^;*tdTomato* mice (*Drp1*^*AgRPKO*^ mice, 14.15 ± 0.926 % of AgRP neurons, n=4; Figure supplement 1b-h) compared to fasted *Drp1*^*+/+*^*-AgRP-cre:ER*^*T2*^;*tdTomato* mice (used as control to visualize AgRP neurons; 52.39 ± 3.71 % of AgRP neurons, n=4, p<0.0001, Figure supplement 1b-h) by immunohistochemistry analysis. No difference in AgRP cell counts were found between control (150.3 ± 4.423 neurons, n=4) and *Drp1*^*AgRPKO*^ mice (146 ± 3.082 neurons, n=4, p=0.4605, Figure supplement 1i).

### Deletion of Drp1 attenuates fasting-induced mitochondrial fission in AgRP neurons

Next, we analyzed mitochondrial morphological changes in AgRP neurons of *Drp1*^*AgRPKO*^ male mice in fed and fasted states. No differences in mitochondrial size (Figure 3a-c), density (Figure 3d), aspect ratio (Figure 3e), and coverage (Figure 3e) were observed between fed and fasted *Drp1*^*AgRPKO*^ male mice, indicating that selective deletion of Drp1 in AgRP neurons prevents fasted-induced mitochondrial fission.

**Figure 3.**
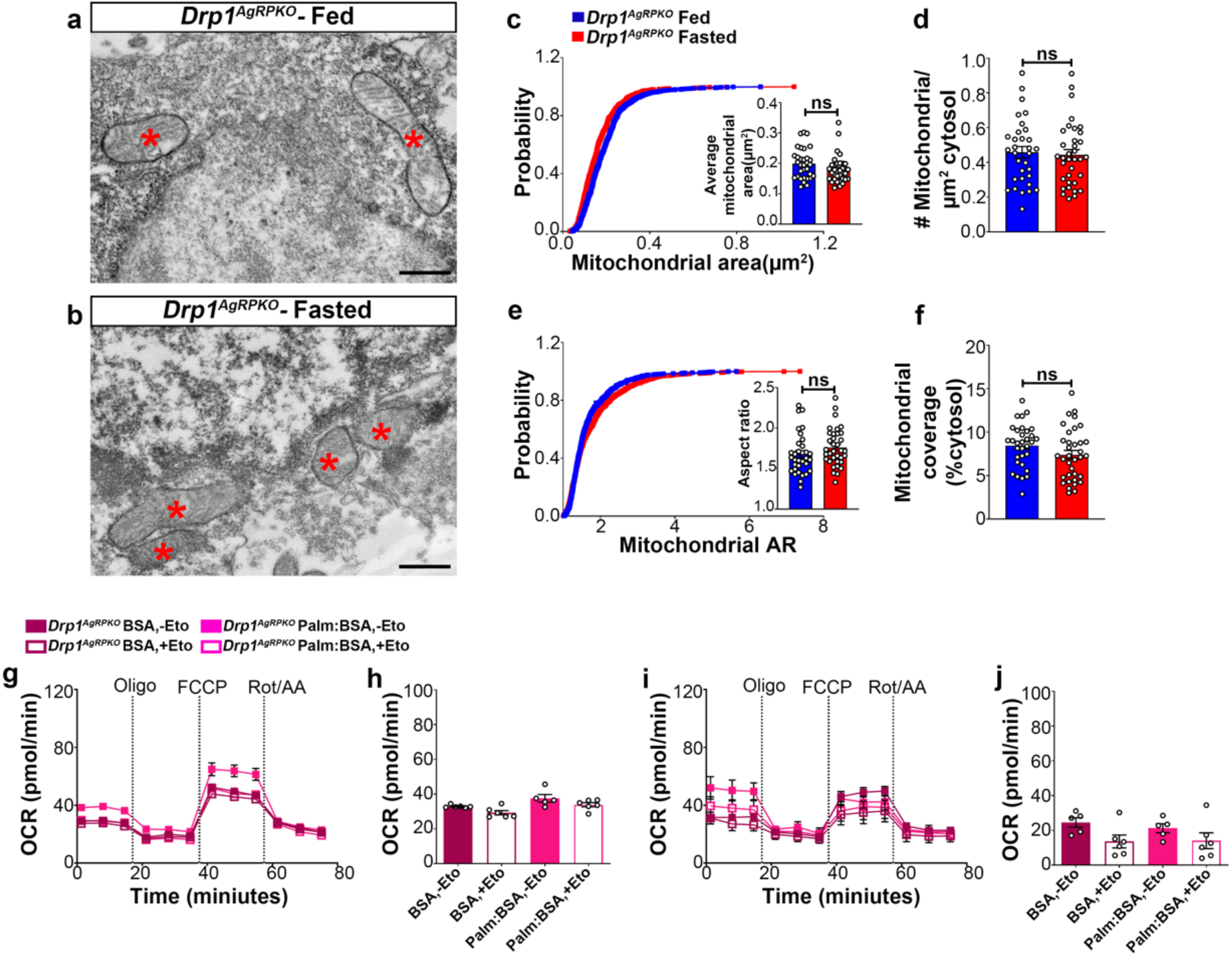
Deletion of DRP1 in AgRP neurons affects fasting-induced mitochondrial fission and mitochondrial respiration. **(a**,**b)** Representative electron micrographs showing mitochondria (asterisks) in an AgRP neuron of the 5-month-old fed *Drp1*^*AgRPKO*^ (a) and the fasted *Drp1*^*AgRPKO*^ male mice (b). Scale bar represents 500 nm. (**c-f)** Cumulative probability distribution of cross-sectional mitochondria area and average mitochondrial area (c), a cumulative probability distribution of mitochondrial aspect ratio and aspect ratio (d), mitochondrial density (e), and mitochondrial coverage (f) in AgRP neurons from fed *Drp1*^*AgRPKO*^ (n=720 mitochondria/32 AgRP neurons/4 mice) and fasted *Drp1*^*AgRPKO*^ male mice (n=746 mitochondria/35 AgRP neurons/4 mice). Data are presented as mean ± SEM. Two-tailed Student’s *t*-test was used for statistical significance. ns=not significant. **(g-h)** Graphs showing OCR (g) and its quantification (h) under 2.5 mM glucose incubation with or without palmitate-BSA (200 μM) and with or without etomoxir (40 μM) in primary hypothalamic neuronal culture of *Drp1*^*AgRPKO*^ mice (n=6-8/ group). Data are presented as mean ± SEM. Two-way ANOVA with Tukey’s post hoc analysis for multiple comparisons was used for statistical significance. **(i-j)** Graphs showing OCR (i) and its quantification (j) under low glucose (0.5 mM) incubation with or without palmitate-BSA (200 μM) and with or without etomoxir (40 μM) in primary hypothalamic neuronal culture of *Drp1*^*AgRPKO*^ mice (n=6-8/ group). Data are presented as mean ± SEM. Two-way ANOVA with Tukey’s post hoc analysis for multiple comparisons was used for statistical significance.

### Deletion of Drp1 in AgRP neurons affects mitochondrial functions

We then determined FA-induced mitochondrial oxygen consumption rate in primary hypothalamic neuronal cell cultures from *Drp1*^*AgRPKO*^ mice. Contrary to our data using control mice (Fig. 2), no difference in FA-induced maximal oxygen consumption rate were observed in high (Figure 3g-h) or low glucose media (Figure 3i-j). In addition, no effects induced by etomoxir incubation were observed in primary hypothalamic neurons derived from *Drp1*^*AgRPKO*^ mice (Figure 3g-j), indicating that Drp1 in the hypothalamic AgRP neurons plays an essential role in regulating FA-induced mitochondrial respiration.

### Inducible and selective deletion of Drp1 in AgRP neurons affect neuronal activation and projection of AgRP neurons in the hypothalamus

To assess the effect of Drp1 deletion on AgRP neuronal function, we then performed and analyzed immunostaining for c-Fos in the hypothalamic arcuate nucleus of *Drp1*^*AgRPKO*^ male mice and controls in fasting state. A significant decrease in immunoreactivity for c-Fos was observed in AgRP neurons of fasted *Drp1*^*AgRPKO*^ male mice (24.2 ± 3.58 % of AgRP neurons, n=5, p=0.0097, Figure 4b,d,f,g) compared to fasted controls (39.14 ± 2.925 % of AgRP neurons, n=6, Figure 4a,c,e,g). No changes in AgRP cell number were observed between control (150.000 ± 5.688 counts, n=6) and *Drp1*^*AgRPKO*^ male mice (150.833 ± 2.753 counts, n=5, p=0.9046, Figure 4h). In agreement with reduced AgRP neuronal activation, a significant reduction in overnight fasting-induced food intake was observed in *Drp1*^*AgRPKO*^ mice (5.155 ± 0.294 g, n=11, p=0.005) compared to controls (6.391 ± 0.290 g, n=11, Figure 4i).

**Figure 4.**
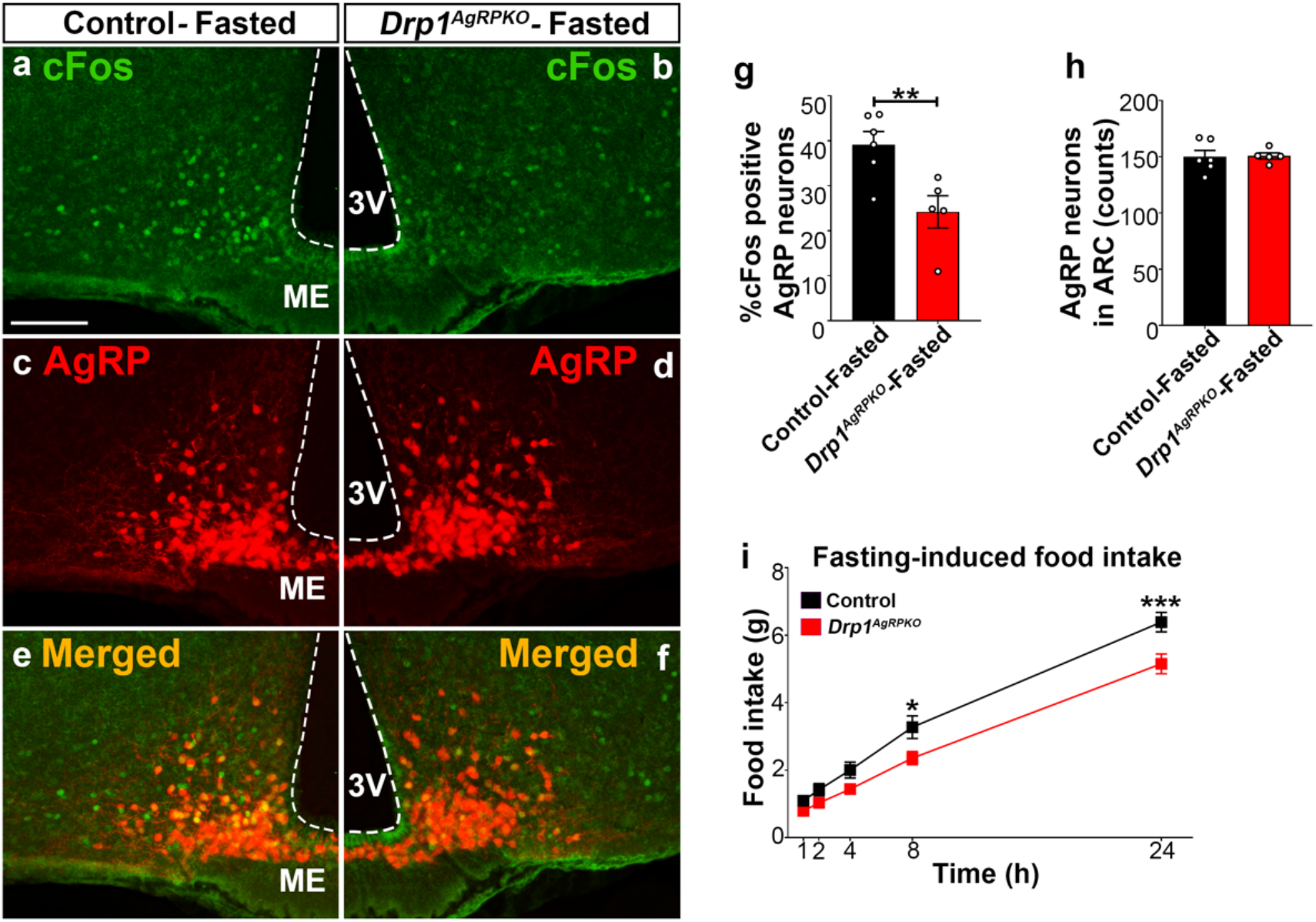
Drp1 deficiency in AgRP neurons affects neuronal activation of the hypothalamic AgRP neurons. **(a-f)** Immunostaining for c-Fos (green, a-b) and *td*Tomato (red, representing AgRP in *Drp1*^*+/+*^*-AgRP-cre:ER*^*T2*^ mice injected with TMX; c-d) and merged (e-f) in the hypothalamic ARC of a fasted male control (a,c,e) and a *Drp1*^*AgRPKO*^ mouse (b,d,f) at 5 months of age. **(g)** Graph showing the percent of c-Fos positive AgRP neurons in fasted control (*Drp1*^*+/+*^*-AgRP-cre:ER*^*T2*^ mice injected with TMX) and *Drp1*^*AgRPKO*^ male mice (n=5-6 mice). **(h)** Graph showing the number of AgRP neurons in the ARC of control and *Drp1*^*AgRPKO*^ mice. Data are presented as mean ± SEM. **P<0.01 by two-tailed Student’s *t*-test. **(i)** Graph showing food intake in male control and *Drp1*^*AgRPKO*^ mice (n=11 mice) after overnight fasting. Data are presented as mean ± SEM. *P<0.05; ***P<0.001 by two-way ANOVA with Tukey’s post hoc analysis for multiple comparisons. 3v=third ventricle; ME=median eminence; ARC=arcuate nucleus.

Furthermore, we analyzed AgRP immunoreactive fibers in target areas of the hypothalamus of fasted *Drp1*^*AgRPKO*^ male mice and controls (Figure 5a-b). Compared to controls, we observed a significant decrease in the ARC AgRP fluorescent intensity (control= 1.000 ± 0.072, n=4; *Drp1*^*AgRPKO*^ mice=0.362 ± 0.070, n=4, p=0.0007, Figure 5c-e) and particle number (control=820.250 ± 34.463 counts, n=4; *Drp1*^*AgRPKO*^ mice=611.25 ± 43.924, n=4, p=0.0096, Figure 5f), in the dorsomedial hypothalamic nucleus (DMH) AgRP fluorescent intensity (relative intensity, control=1.000 ± 0.099, n=4; *Drp1*^*AgRPKO*^ mice=0.262 ± 0.019, n=4, p=0.0003, Figure 5g-i) and particle number (control =803.688 ± 27.808 counts, n=4; *Drp1*^*AgRPKO*^ mice=385.500 ± 17.093, n=4, p<0.0001, Figure 5j) and in the paraventricular nucleus (PVN) AgRP fluorescent intensity (relative intensity, control=1.000 ± 0.093, n=4; *Drp1*^*AgRPKO*^ mice=0.308 ± 0.038, n=4, p=0.0005, Figure 5k-m) and particle number (control=877.438 ± 35.425 counts, n=4; *Drp1*^*AgRPKO*^ mice=655.750 ± 26.976, n=4, p=0.0025, Figure 5n). Similar results were also observed, for example, in the PVN of *Drp1*^*AgRPKO*^ female mice compared to controls (Figure supplement 2).

**Figure 5.**
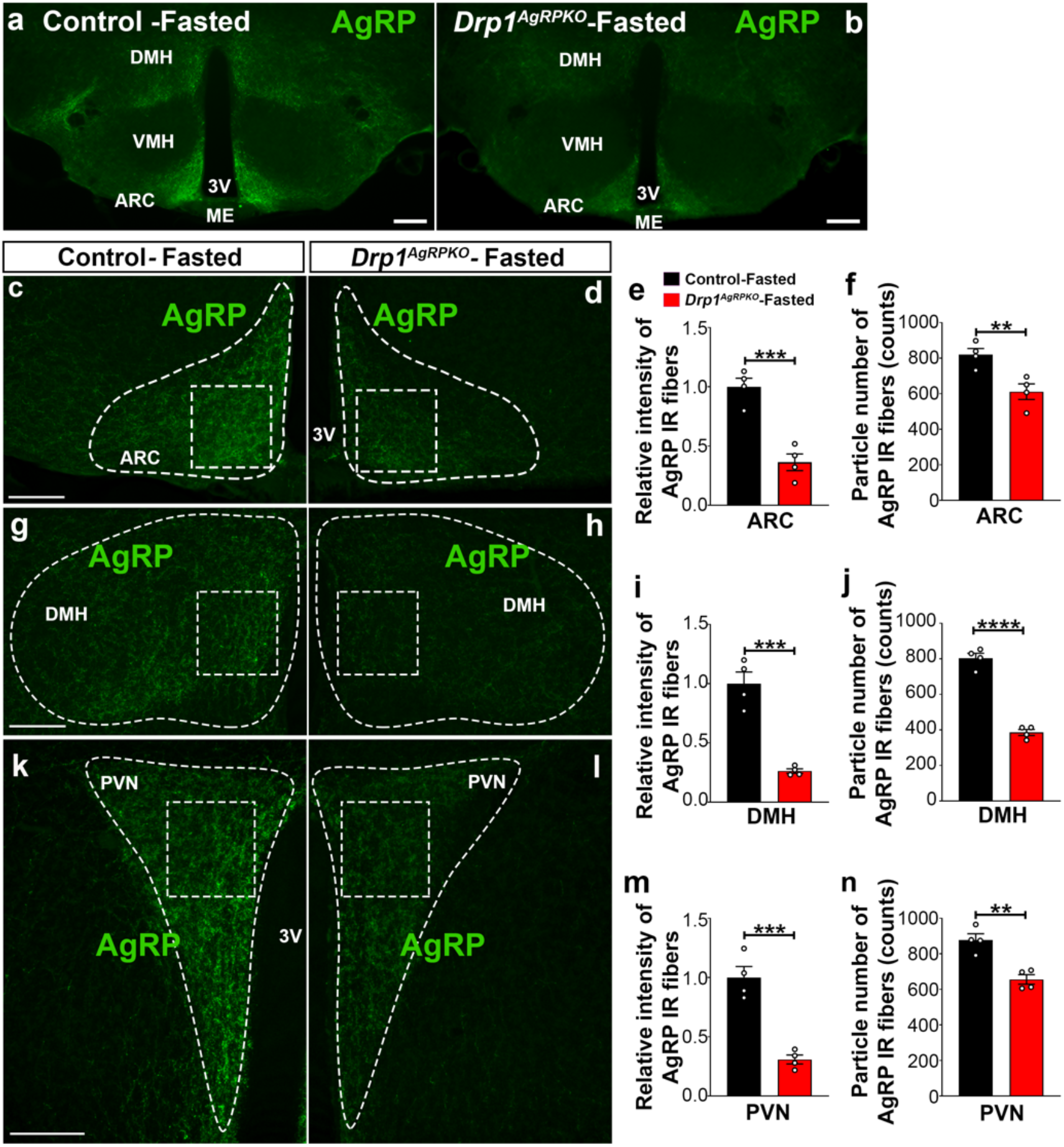
AgRP-selective Drp1 deficiency affects AgRP projections within the hypothalamus. **(a-b)** Immunostaining for AgRP (green) in the hypothalamus of a fasted male control (a) and a fasted *Drp1*^*AgRPKO*^ mouse (b) at 5 months of age. **(c-d)** Immunostaining for AgRP in the ARC of a fasted control (c) and a fasted *Drp1*^*AgRPKO*^ mouse (d). Dashed lines delineate the ARC. Squared area represents the area used for for analysis. (**e-f)** Graphs showing the quantification of relative intensity (e) and particle number (f) of AgRP fibers in the ARC of fasted control and *Drp1*^*AgRPKO*^ male mice (n=4 mice). **(g-h)** Immunostaining for AgRP in DMH of a fasted control (g) and a fasted *Drp1*^*AgRPKO*^ mouse (h). Dashed lines delineate the DMH. Squared area represents the area used for analysis. **(i-j)** Graphs showing the quantification of relative intensity (i) and particle number (j) of AgRP fibers in DMH of fasted control and *Drp1*^*AgRPKO*^ male mice (n=4 mice). **(k-l)** Immunostaining for AgRP in the PVN of a fasted control (k) and a fasted *Drp1*^*AgRPKO*^ mouse (l). Dashed lines delineate the PVN. Squared area represents the area used for analysis. **(m-n)** Graphs showing the quantification of relative intensity (m) and particle number (n) of AgRP fibers in the PVN of fasted control and *Drp1*^*AgRPKO*^ male mice (n=4 mice). Scale bar represents 100 µm (c,g,k) and 200 µm (a-b). All data are presented as mean ± SEM. **P<0.01; ***P<0.001; ****P<0.0001 by two-tailed Student’s *t*-test. 3v=third ventricle; ME=median eminence; DMH=dorsomedial hypothalamus; VMH=ventromedial hypothalamus; PVN=paraventricular hypothalamus; ARC=arcuate nucleus.

### Deletion of Drp1 in AgRP neurons affects POMC and paraventricular neuronal activation

Next, we analyzed immunostaining for c-Fos in POMC neurons of *Drp1*^*AgRPKO*^ male mice and their controls (Figure supplement 3). A significant increase in POMC cells immunoreactive for c-Fos was observed in *Drp1*^*AgRPKO*^ mice (33.683 ± 2.050 % of POMC neurons, n=4, p=0.0032) compared to controls (22.169 ± 1.297 % of POMC neurons, n=4, Figure supplement 3a-g). No changes in POMC cell number were observed between control and *Drp1*^*AgRPKO*^ mice (Figure supplement 3h).

We then analyzed α-MSH fiber immunostaining in the PVN of fasted *Drp1*^*AgRPKO*^ male mice and their controls (Figure supplement 3i-j). Significant increases in relative intensity (control=1.000 ± 0.164 counts, n=4; *Drp1*^*AgRPKO*^ mice=6.195 ± 0.494, n=5, p<0.0001, Figure supplement 3k) and particle number (control=157.625 ± 13.488 counts, n=4; *Drp1*^*AgRPKO*^ mice=393.400 ± 19.290 counts, n=5, p<0.0001, Figure supplement 3l) of α-MSH fibers were observed in the PVN of *Drp1*^*AgRPKO*^ mice compared to their controls.

In agreement with a reduced AgRP- and an increased POMC neuronal activation, we observed a significant increase in c-Fos immunopositive cells in the PVN of fasted *Drp1*^*AgRPKO*^ mice (72.600 ± 9.092 counts, n=5, p=0.0028) compared to their controls (22.750 ± 4.535 counts, n=4, Figure supplement 3m-o).

### Deletion of Drp1 in AgRP neurons alters energy metabolism

To determine the physiological role of Drp1 in AgRP neurons in regulating energy metabolism, we assessed the metabolic phenotype of male and female *Drp1*^*AgRPKO*^ mice and their controls. Before starting tamoxifen (TMX) injections at 5 weeks of age, no significant differences in body weight were observed between controls and *Drp1*^*AgRPKO*^ mice in male (Body weight=*Drp1*^*+/+*^*-AgRP-cre:ER*^*T2*^-TMX=18.533 ± 0.390, n=18; *Drp1*^*fl/fl*^*-AgRP-cre:ER*^*T2*^*-*TMX=18.176 ± 0.512, n=17; *Drp1*^*fl/fl*^*-AgRP-cre:ER*^*T2*^-Corn oil=18.830 ± 0.308, n=10; p=0.8649 for *Drp1*^*+/+*^*-AgRP-cre:ER*^*T2*^-TMX versus *Drp1*^*fl/fl*^*-AgRP-cre:ER*^*T2*^*-*TMX ; p=0.9289 for *Drp1*^*+/+*^*-AgRP-cre:ER*^*T2*^-TMX versus *Drp1*^*fl/fl*^*-AgRP-cre:ER*^*T2*^-Corn oil; p=0.7045 for *Drp1*^*fl/fl*^*-AgRP-cre:ER*^*T2*^*-*TMX versus *Drp1*^*fl/fl*^*-AgRP-cre:ER*^*T2*^-Corn oil, Figure 6a).

**Figure 6.**
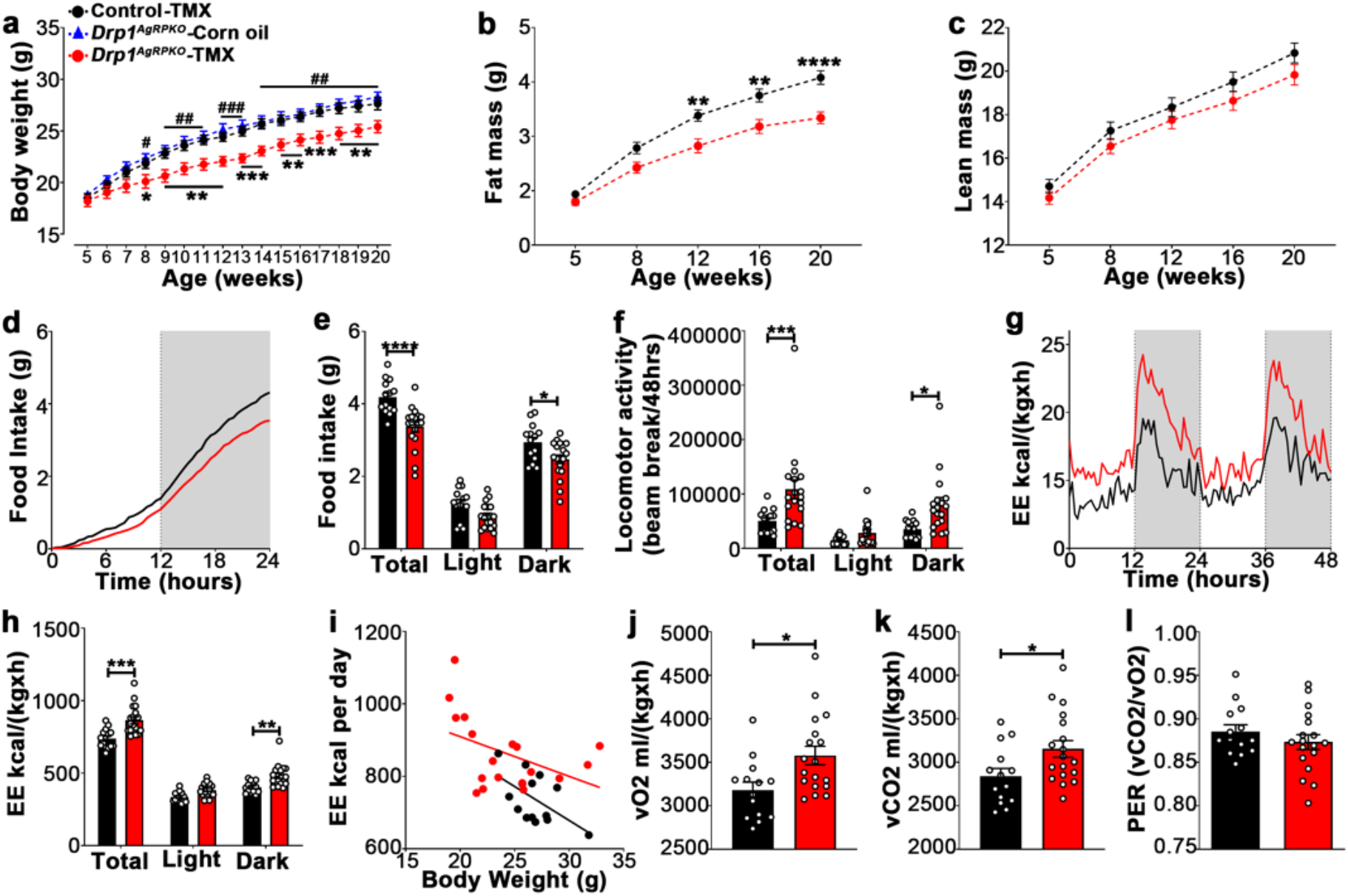
Deletion of DRP1 in AgRP neurons affects metabolic phenotype in male mice. **(a)** Graph showing body weight of *Drp1*^*+/+*^*-AgRP-cre:ER*^*T2*^ mice injected with tamoxifen (n=18 mice), *Drp1*^*fl/fl*^*-AgRP-cre:ER*^*T2*^ mice injected with corn oil (n=10 mice) as control groups, and *Drp1*^*fl/fl*^*-AgRP-cre:ER*^*T2*^ mice injected with tamoxifen (n=17 mice). Data are presented as mean ± SEM. *P<0.05; **P<0.01; ***P<0.001 for *Drp1*^*+/+*^*-AgRP-cre:ER*^*T2*^-TMX versus *Drp1*^*fl/fl*^*-AgRP-cre:ER*^*T2*^*-*TMX; ^#^P<0.05; ^##^P<0.01; ^###^P<0.001 for *Drp1*^*fl/fl*^*-AgRP-cre:ER*^*T2*^-Corn oil versus *Drp1*^*fl/fl*^*-AgRP-cre:ER*^*T2*^-TMX by two-way ANOVA with Tukey’s post hoc analysis for multiple comparisons. **(b-c)** Graphs showing fat mass (b), and lean mass (c) of *Drp1*^*+/+*^*-AgRP-cre:ER*^*T2*^ mice (n=20 mice) and *Drp1*^*fl/fl*^*-AgRP-cre:ER*^*T2*^ mice (n=22 mice). Data are presented as mean ± SEM. **P<0.01; ****P<0.0001 by two-way ANOVA with Tukey’s post hoc analysis for multiple comparisons. **(d-e)** Graphs showing cumulative 24 hours food intake in 4 months old control (n=14) and *Drp1*^*AgRPKO*^ male mice (n=18) (d), and results of food intake as total in the 24 h cycle and in the dark and light phases of the cycle (e; average of 3 days). Gray area represents dark phases. Data are presented as mean ± SEM. *P<0.05; ****P<0.0001 by two-way ANOVA with Tukey’s post hoc analysis for multiple comparisons. **(f-l)** Graphs showing locomotor activity (f), energy expenditure (g-i), consumed O_2_ (j), produced CO_2_ (k), and the respiratory exchange ratio (RER) (l) in 4 months old control (n=14) and *Drp1*^*AgRPKO*^ male mice (n=18). Data are presented as mean ± SEM. *P<0.05; **P<0.01; ***P<0.001 by two-way ANOVA with Tukey’s post hoc analysis for multiple comparisons. p=0.5203 by linear regression analysis (i). *P<0.05 by two-tailed Student’s *t*-test (j-k).

A significant decrease in body weight in *Drp1*^*AgRPKO*^ male mice compared to controls was observed first at 8 weeks of age, 3 weeks after the start of TMX treatment (Figure 6a) and was maintained through the end of the study when the mice were 20 weeks old (Figure 6a; n=22 per group).

The decrease in body weight was associated with a significant reduction in fat mass (Figure 6b; n=22, p<0.0001) while no significant difference in lean mass was observed (Figure 6c; n=22, p=0.3421) compared to control mice.

*Drp1*^*AgRPKO*^ mice showed significantly lower 24 h food intake (control=4.181 ± 0.124 g, n=14; *Drp1*^*AgRPKO*^ mice=3.366 ± 0.139 g, n=18, p<0.0001, total in Figure 6d-e), specifically during the dark period compared to controls (control=2.937 ± 0.137 g, n=14; *Drp1*^*AgRPKO*^ mice=2.471 ± 0.120 g, n=18, p=0.0236, Dark in Figure 6e). A significant increase in locomotor activity was observed in *Drp1*^*AgRPKO*^ mice (control= 50122.750 ± 5919.799 beam brake counts, n=14; *Drp1*^*AgRPKO*^ mice=108400.917 ± 17432.685 beam brake counts, n=18, p=0.0008, Figure 6f), specifically during the dark period compared to controls (control=35556.429 ± 4272.846 counts, n=14; *Drp1*^*AgRPKO*^ mice=80081.806 ± 13085.949 counts, n=18, p=0.0136, Figure 6f).

The difference in body weight and composition were also associated with significantly increased energy expenditure (control=738.227 ± 18.025, n=14; *Drp1*^*AgRPKO*^ mice =864.916 ± 23.597, n=18, p<0.0001, Figure 6g-i), increased O2 consumption (control=3179 ± 92.26, n=14; *Drp1*^*AgRPKO*^ mice=3578 ± 106.6, n=18, p=0.0104, Figure 6j) and CO2 production (control=2840 ± 89.27, n=14; *Drp1*^*AgRPKO*^ mice=3154 ± 95.78, n=18, p=0.0262, Figure 6k) in *Drp1*^*AgRPKO*^ mice compared to control mice, while no significant differences in the respiration exchange rate (RER) were observed between *Drp1*^*AgRPKO*^ mice (0.8734 ± 0.00852, n=18, p= 0.3133) and control mice (0.8855 ± 0.007676, n=14, Figure 6l). Similar to males, female *Drp1*^*AgRPKO*^ mice showed significant differences in body weight, composition, feeding, energy expenditure and locomotor activity compared to controls (Figure supplement 4).

### Deletion of Drp1 in AgRP neurons results in increased brown adipose tissue thermogenesis

BAT thermogenesis is a critical component of the homeostatic energy balance to maintain body temperature (Morrison and Madden, 2014). We then, examined whether deletion of Drp1 in AgRP neurons affects body temperature. We found that BAT temperature was significantly increased in *Drp1*^*AgRPKO*^ mice compared to control mice (control=33.388 ± 0.223 °C, n=8; *Drp1*^*AgRPKO*^ mice=35.038 ± 0.318 °C, n=8, p=0.0008, Figure supplement 5a-c). Rectal temperature was also significantly increased in *Drp1*^*AgRPKO*^ mice (37.300 ± 0.105 °C, n=8) compared to controls (36.654 ± 0.226 °C, n=8; p=0.0283, Figure supplement 5d). Similar to males, rectal temperature of female *Drp1*^*AgRPKO*^ mice (37.181 ± 0.085 °C, n=7) was significantly greater than that of control (36.652 ± 0.142 °C, n=9; p=0.0102; Figure supplement 5e).

### Ghrelin-induced hyperphagia and AgRP activation is dependent on Drp1

Ghrelin, a gut-derived hormone secreted during food deprivation, promotes feeding behavior through NPY/AgRP neurons (Andrews et al., 2008). We found a significant decrease in c-Fos immunoreactivity in AgRP neurons of ghrelin-treated *Drp1*^*AgRPKO*^ mice (30.22 ± 4.652 % of AgRP neurons, n=5, p=0.0001) compared to controls (75.35 ± 3.464 % of AgRP neurons, n=4, Figure 7a-g). No difference in the number of AgRP neurons in the ARC was observed between the 2 groups (Figure 7h). In agreement with that, ghrelin-induced hyperphagia was not observed in *Drp1*^*AgRPKO*^ mice compared to their controls (Figure 7i). Next, we performed patch-clamp whole-cell electrophysiological recordings in slices from *Drp1*^*AgRPKO*^ mice and controls. In support of the c-Fos data, ghrelin significantly increased membrane potential (resting=−46.644 ± 0.502 mV, n=20; ghrelin=−43.757 ± 0.678 mV, n=20, p=0.0102; Figure 7j,l) and relative firing activity (resting=100.000 ± 14.584, n=20; ghrelin=186.894 ± 20.266, n=20, p=0.0041; Figure 7k-l) of AgRP neurons in control mice, while ghrelin-induced excitation of AgRP neurons was significantly attenuated in *Drp1*^*AgRPKO*^ mice compared to controls (membrane potential, resting=−46.104 ± 0.577 mV, n=19; ghrelin=−45.677 ± 0.644 mV, n=19, p=0.9962, Figure 7j,l; relative firing activity, resting=100.000 ± 15.106, n=19; ghrelin=119.985 ± 16.502, n=19, p=0.9644, Figure 7k-l). Note that no differences in the levels of total (Figure supplement 6a) and active form ghrelin (Figure supplement 6b) in the fed or fasted states between male control and *Drp1*^*AgRPKO*^ mice were observed. Together, these data suggest that Drp1-mediated mitochondrial fission plays a critical role in regulating ghrelin-triggered AgRP neuronal activity and hyperphagia.

**Figure 7.**
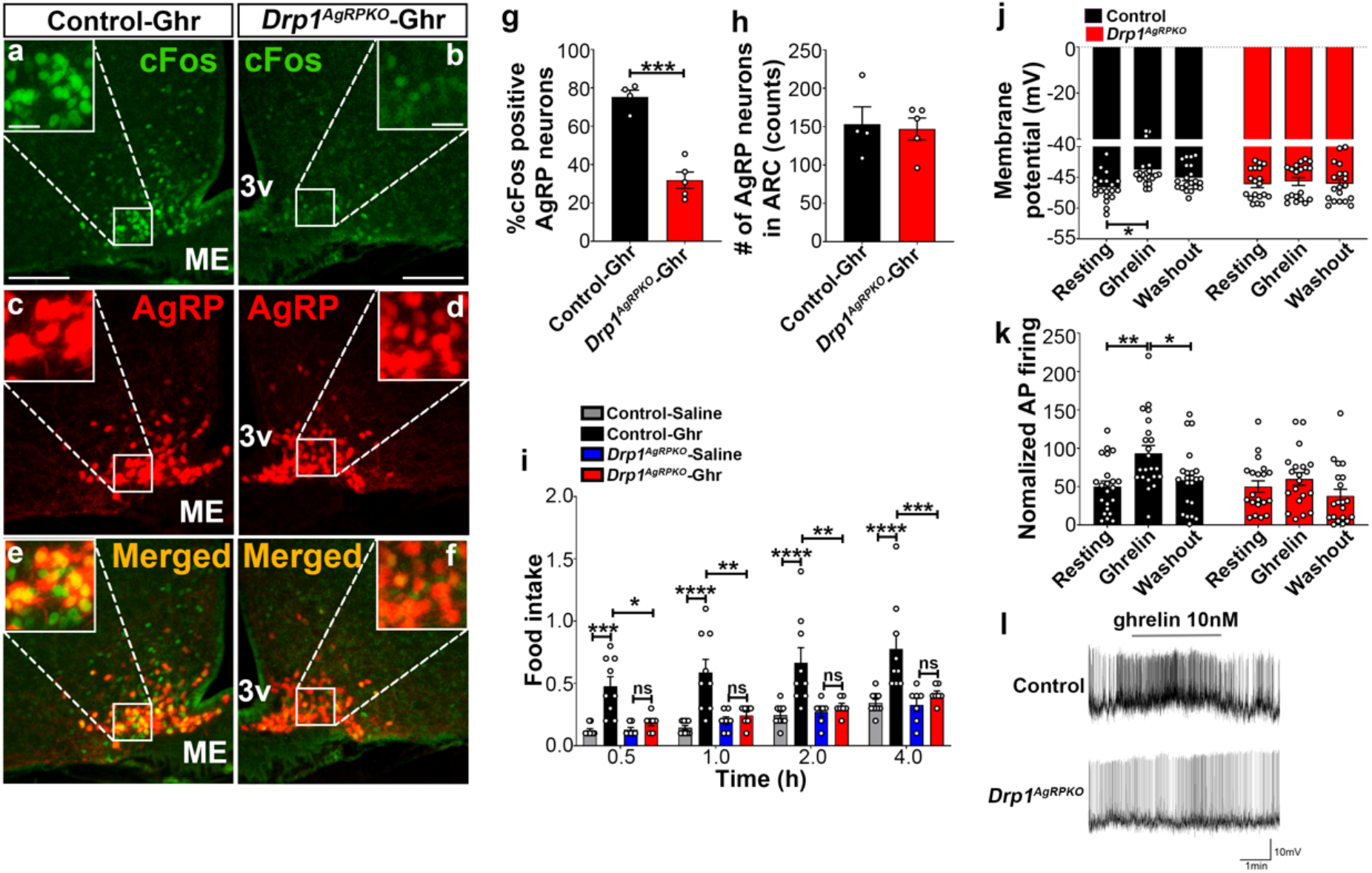
Deletion of Drp1 in AgRP neurons attenuates ghrelin-induced neuronal activation and feeding. **(a-f)** Immunostaining for c-Fos (green, a-b) and *td*Tomato (red, representing AgRP, c-d) and merged (e-f) in the ARC of a ghrelin-injected male control (a,c,e) anda *Drp1*^*AgRPKO*^ mouse (b,d,f). **(g)** Graph showing the quantification of c-Fos expression in AgRP neurons of ghrelin-injected control and *Drp1*^*AgRPKO*^ mice (n=4-5 mice). **(h)** Graph showing the number of AgRP neurons of ghrelin-injected control and *Drp1*^*AgRPKO*^ mice in the hypothalamic ARC (n=4-5 mice). Data are presented as mean ± SEM. ***=P<0.001 by two-tailed Student’s *t*-test. **(i)** Food intake in 4 months old control and *Drp1*^*AgRPKO*^ female mice (n=7-9 mice/group) after either saline or ghrelin injection. *P<0.05; **P<0.01; ***P<0.001; ****P<0.0001; ns=not significant; Two-way ANOVA with Tukey’s post hoc analysis for multiple comparisons was performed. **(j)** Graph showing the membrane potential in AgRP neurons of 3 months old control (n=20 cells/10 mice) and *Drp1*^*AgRPKO*^ male mice (n=19 cells/10 mice) in response to ghrelin. **(k)** Graph showing normalized firing rate in AgRP neurons of 3 months old control (n=20 cells/10 mice) and *Drp1*^*AgRPKO*^ male mice (n=19 cells/10 mice) in response to ghrelin. Data are presented as mean ± SEM. **=P<0.01 for artificial CSF-treated control versus ghrelin-treated control; *=P<0.05 for ghrelin-treated control versus washed out control by two-way ANOVA with Tukey’s post hoc analysis for multiple comparisons. ns=not significant. **(l)** Representative tracers of AgRP neurons from a control and a *Drp1*^*AgRPKO*^ mouse. Scale bar represents 100 µm. Scale bar in high magnification image represents 20 µm. 3v=third ventricle; ME=median eminence

## Discussion

Our findings revealed a crucial role of AgRP neuronal mitochondrial fission in the regulation of hypothalamic feeding control. First, we found that activated AgRP neurons have decreased mitochondrial size accompanied by an increase in mitochondria number suggesting a mitochondrial fission process. In agreement with this, we found that Drp1 mRNA levels and Drp1 activation (Liesa et al., 2009) are significantly increased in AgRP neurons of fasted compared to fed mice. These data were associated with a significant increase in FA-induced mitochondrial respiration when low glucose levels (similar to fasting) were present compared to high glucose levels. To determine the physiological relevance of mitochondrial fission in AgRP neurons, we generated a mouse model lacking Drp1 in AgRP neurons (*Drp1*^*AgRPKO*^). We found that *Drp1*^*AgRPKO*^ mice, in which fasting did not induced mitochondrial fission and changes in mitochondrial function, had significant decreases in body weight, composition, and feeding that were accompanied by increases in locomotion and energy expenditure. Finally, *Drp1*^*AgRPKO*^ mice also showed attenuated ghrelin-induced hyperphagia and neuronal activity of AgRP neurons. Altogether, these data revealed that DRP1-driven mitochondrial fission in AgRP neurons is an innate adaptive process enabling these neurons to respond to the changing metabolic environment.

Mitochondria are energy-producing organelles fundamental in support of cellular functions. Mitochondria are highly dynamic organelles able not only to move within the cell to sites where their function is required, but they are also able to fuse (mitochondrial fusion) and divide (mitochondrial fission) in order to maintain proper cellular function.

Mitochondrial fusion and fission are highly regulated processes. Several proteins are involved in these events, including MFN1 and MFN2 and OPA1 for mitochondrial fusion, and, Fis1, Mff and Drp1 for mitochondrial fission (Pozo Devoto and Falzone, 2017). Mitochondrial dynamics through fusion and fission processes are also important in maintaining mitochondrial quality control in order to maintain optimal mitochondrial bioenergetic functions (Twig et al., 2008). Our data indicate that mitochondrial dynamics and specifically mitochondrial fission plays an important role in sensing changes of nutrients availability in AgRP neurons. First, we observed that incubation of primary hypothalamic neurons with palmitic acid induced a significant increase in mitochondrial respiration when glucose levels were low, mimicking fasting. Fasting induced increased AgRP neuronal activation and increased mitochondrial fission. However, when Drp1-induced mitochondrial fission in AgRP neurons was abolished, palmitic acid-induced mitochondrial respiration was diminished. In association with these, ghrelin-triggered changes in membrane potential and firing frequency of AgRP neurons were significantly attenuated in *Drp1*^*AgRPKO*^ mice, leading to failure in inducing hyperphagia. In line with our results, Dietrich et al., (2013) have shown that in mice with AgRP-selective deletion of *Mfn1* and *Mfn2*, mediators of mitochondrial fusion process, neuronal firing frequency was impaired in DIO mice. The impairment of AgRP neuronal activation was reversed by increasing intracellular ATP levels (Dietrich et al., 2013), indicating that the impaired AgRP neuronal firing frequency is likely due to low intracellular ATP levels.

Overall, our data unmask that mitochondrial fission in hypothalamic AgRP neurons is a fundamental mechanism that allows these neurons to sense and respond to changes circulating signals, including hormones such as ghrelin and nutrients, such as glucose and palmitic acid, in the regulation of feeding and energy metabolism.

## Methods and materials

### Animals

All animal care and experimental procedures done in this study were approved by the Yale University Institutional Animal Care and Use Committee. All mice were housed in a temperature-controlled environment (22°C-24°C) with a 12-hr light and 12-hr dark (19.00–07.00) photoperiod. Animals were provided standard chow diet (SD) (2018; 18% calories from fat; Harlan Teklad) and water *ad libitum* unless otherwise stated. All mice studied were of the same background.

### Generation of experimental mice with inducible deletion of Drp1 specifically in AgRP neurons

We used the inducible Cre/loxP technology to generate mice in which Drp1 was selectively ablated in AgRP neurons (*Drp1*^*AgRPKO*^ mice). First, mice expressing a tamoxifen-inducible Cre recombinase (*CreER*^*T2*^) in cells expressing AgRP (*AgRP-cre:ER*^*T2*^) were crossed with Rosa26-lox-stop-lox-tdTomato (*Ai14*; cre-recombinase-dependent expression) mice (Ai14 reporter mice) to label AgRP-expressing cells. *AgRP-cre:ER*^*T2*^; *Rosa26-lox-stop-lox-tdTomato* (*AgRP-cre:ER*^*T2*^; *tdTomato*) mice have AgRP-expressing cells with the expression of *td*Tomato by tamoxifen administration. No observation of AgRP-*td*Tomato expression was found in the absence of tamoxifen administration, indicating that recombination was strictly dependent upon tamoxifen-induced Cre recombinase activation. The mice with *AgRP-cre:ER*^*T2*^; *tdTomato* were then crossed with mice harboring conditional alleles *Drp1* floxed (*Drp1*^*fl/fl*^; (Kageyama et al., 2014)) to generated mice with inducible deletion of Drp1 specifically in AgRP neurons (*Drp1*^*AgRPKO*^ mice). *Drp1*^*fl/fl*^*-AgRP-cre:ER*^*T2*^;*tdTomato* mice injected with corn oil and *Drp1*^*+/+*^*-AgRP-cre:ER*^*T2*^;*tdTomato* mice injected with tamoxifen (TMX) were used as controls. *Drp1*^*fl/fl*^*-AgRP-cre:ER*^*T2*^;*tdTomato* mice were injected intraperitoneally (i.p.) with tamoxifen (0.10 mg/g BW for every 3 days with 5 times fasting) starting at 5 weeks of age to induce mature-onset deletion of Drp1 in AgRP neurons of *Drp1*^*AgRPKO*^ mice, and *Drp1*^*+/+*^*-AgRP-cre:ER*^*T2*^;*tdTomato* mice were injected with tamoxifen and *Drp1*^*fl/fl*^*-AgRP-cre:ER*^*T2*^;*tdTomato* were mice injected with corn oil as control groups. Because we found no differences between these two control groups, the majority of the experiments were performed using *Drp1*^*+/+*^*-AgRP-cre:ER*^*T2*^;*tdTomato* mice injected with tamoxifen (to label AgRP neurons with *td*Tomato expression) as a control group, unless otherwise stated.

### Ribotag assays

We performed transcriptomic profiling by using ribosomal tagging strategy to analyze AgRP neurons-specific mRNA expression in vivo. To avoid the potential disadvantage that the embryonic POMC-expressing progenitor neurons differentiate into AgRP-expressing neurons, we crossed *AgRP-cre:ER*^*T2*^ mice with *Rpl22* floxed (RiboTag) mice to eventually generate *AgRP-cre:ER*^*T2*^; *RiboTag* mice, expressing a hemagglutinin A (HA)-tagged ribosomal protein in the AgRP neurons upon tamoxifen injection. Eleven to twelve-week-old mice (one month after the last tamoxifen injection) were used. After mice were anesthetized with isoflurane and decapitated, the brains were rapidly dissected out. To collect mediobasal hypothalamus (MBH) containing the Arc, brain tissues were sectioned in two millimeter thick coronal sections containing MBH in a brain matrix. MBH Arc samples were collected under a stereomicroscope according to the brain atlas for appropriate regions and preventing differences in tissue weight. Three animals were pooled for each N. The MBH ARC samples from *AgRP-cre:ER*^*T2*^; *RiboTag* mice were homogenized by supplemented homogenization buffer (HB-S: 50 mM Tris, pH 7.4, 100 mM KCl, 12 mM MgCl2, and 1 % NP-40 supplemented with 1 mM DTT, 1 mg/ml heparin, 100 µg/ml cycloheximide, 200 U/ml RNasin Ribonuclease inhibitor, and protease inhibitor cocktail). Samples were then centrifuged at 10,000 rpm for 10 minutes at 4°C. Then, 50 µl of each supernatant was transferred to a new tube serving as input fraction (containing all mRNAs). To isolate polyribosomes, we performed immunoprecipitation of ribosome-bound mRNAs in AgRP neurons. by utilizing anti-HA antibody (5 µl/ sample; Cat#901513, Biolegend).

RNA was extracted using Qiagen RNeasy® Plus Micro Kit (Cat# 74034, Qiagen) according to the protocol supplied by the manufacturer. cDNA was synthesized using High Capacity cDNA Reverse transcription Kit (Cat# 4368814, Thermo Fisher Scientific). qRT-PCR experiment was performed by Taqman Gene Expression Assay primers (Thermo Fisher Scientific) in triplicates using LightCycler 480 Real-Time PCR System (Roche Diagnostics, Mannheim, Germany). All genes were normalized to β-actin. The calculation of average Cp values and resulting expression ratios for each target gene was performed using the Roche LightCycler 480 software. The following primers were utilized: dnm1l, Mm01342903_m1; AgRP, Mm00475829_g1; Npy, Mm01410146_m1; Pomc, Mm00435874_m1; NR5A1, Mm00446826_m1; Gapdh, Mm99999915_g1; β-actin, Mm02619580.

### Metabolic assays

Four months old mice were acclimated in metabolic chambers (TSE System-Core Metabolic Phenotyping Center, Yale University) for 3 days before the start of the recordings. Mice were continuously recorded for 2 days, with the following measurements taken every 30 min: food intake, locomotor activity (in the xy- and z-axes), and gas exchange (O_2_ and CO_2_; The TSE LabMaster System). Energy expenditure was calculated according to the manufacturer’s guidelines (PhenoMaster Software, TSE System). The respiratory quotient was estimated by calculating the ratio of CO_2_ production to O_2_ consumption. Values were adjusted by body weight to the power of 0.75 (kg−0.75) where mentioned. Body composition was measured in vivo by MRI (EchoMRI, Echo Medical Systems, Houston, TX). Body core temperature was measured using a thermocouple rectal probe and thermometer (Physitemp instruments). Rectal temperature was measured repeated three times, and the average was calculated. The temperature of the surface overlying BAT was measured using infrared thermography images (FLIR C2, FLIR Thermal Imaging System). The infrared thermography images were taken at least three times and analyzed using FLIR Tools (FLIR Thermal Imaging System).

### Phosphorylated-Drp1 immunostaining

Five months old mice were deeply anesthetized and transcardially perfused with 0.9% saline containing heparin (10 mg/L), followed by fresh fixative of 4% paraformaldehyde in phosphate buffer (0.1 M PB, pH 7.4) as previously described (Andrews et al., 2008, Diano et al., 2011, Toda et al., 2016). Brains were post-fixed overnight at 4°C and sliced to a thickness of 50 µm using a vibratome (#11000, PELCO easySlicer, TED PELLA Inc.) and coronal brain sections containing the hypothalamic arcuate nucleus (ARC) were selected under the stereomicroscope (Stemi DV4, Carl Zeiss Microimaging Inc.). After several washes with 0.1 M PB, brain sections were preincubated with 0.2% triton X-100 (Sigma-Aldrich) and 2% normal goat serum in 0.1 M PB for 30 min to permeabilize tissue and cells. Brain sections were incubated with rabbit anti-phosphorylated-Drp1 (Ser-616) antibody (diluted 1:500 in 0.1 M PB, #4494, Cell Signaling) overnight at room temperature (RT). The following day, brain sections were washed and incubated with a biotinylated goat anti-rabbit IgG (diluted 1:200 in 0.1M PB, BA-1000, Vector Laboratories) for 2 hr at RT. Sections were then washed and incubated in streptavidin-conjugated Alexa Fluor® 488 (diluted 1:2000 in 0.1 M PB, A21370, Life Technologies) for 2 hr at RT. No staining was performed to visualize AgRP neurons since mice were expressing *td*Tomato in this neuronal population, which is per se fluorescent. After several washes with 0.1 M PB, brain sections were mounted on glass slides and coverslipped with a drop of Vectashield mounting medium (H-1000, Vector Laboratories). The coverslip was sealed with nail polish to prevent drying and movement under the microscope. All slides were stored in the dark at 4°C.

### c-Fos immunostaining

Five months old mice were deeply anesthetized and transcardially perfused as described above. Immunofluorescent staining was performed using rabbit anti-c-Fos antibody (diluted 1:2,000 in 0.1 M PB, sc-52, Santa Cruz Biotechnology) overnight at RT. The following day, brain sections were washed and incubated with a biotinylated goat anti-rabbit IgG (diluted 1:200 in 0.1M PB, BA-1000, Vector Laboratories) for 2 hr at RT. Sections were then washed and incubated in streptavidin-conjugated Alexa Fluor® 488 (diluted 1:2000 in 0.1 M PB, A21370, Life Technologies) for 2 hr at RT. No staining was performed to visualize AgRP neurons since mice were expressing *td*Tomato in this neuronal population, which is per se fluorescent. For POMC immunofluorescence, brain sections were then incubated with rabbit anti-POMC antibody (diluted 1:2,000 in 0.1 M PB, H-029-30, Phoenix Pharmaceuticals). The following day, sections were washed and incubated with anti-rabbit Alexa Fluor® 594 (diluted 1:500 in 0.1 M PB, A11037, Life Technologies) for 2 hr at RT. After several washes with 0.1 M PB, brain sections were mounted on glass slides and coverslipped with a drop of vectashield mounting medium (H-1000, Vector Laboratories) and analyzed with a fluorescence microscope.

### AgRP and α-MSH fiber immunostaining

Five months old mice were deeply anesthetized and transcardially perfused as described above. Brain sections containing the hypothalamic paraventricular nucleus (PVN) were selected under the stereomicroscope. Immunofluorescence staining was performed using sheep anti-α-MSH antibody (diluted 1:1,000 in 0.1 M PB, ab5087, Millipore Sigma), and rabbit anti-AgRP antibody (diluted 1:1,000 in 0.1 M PB, H-003-57, Phoenix Pharmaceuticals, Inc) overnight at RT. The following day, brain sections were washed and incubated with anti-sheep Alexa Fluor® 488 (diluted 1:1,000 in 0.1M PB, A11015, Life technologies), and anti-rabbit Alexa Fluor® 488 (diluted 1:1,000 in 0.1M PB, A21206, Life Technologies) for 2 hr at RT. After several washes with 0.1 M PB, brain sections were mounted on glass slides and coverslipped with a drop of vectashield mounting medium and analyzed with a fluorescence microscope.

### Fluorescent image capture and analyses

Five months old mice were deeply anesthetized and transcardially perfused as described above. Fluorescent images were captured with Fluorescence Microscope (Model BZ-X710, KEYENCE). For all immunohistochemistry (IHC) analyses, coronal brain sections were anatomically matched (ARC: between −1.46 and −1.94 mm from bregma, PVN: −0.82 and −0.94 mm from bregma) with the mouse brain atlas (Franklin and Paxinos, 2019). Both sides of the bilateral brain region (ARC and PVN) were analyzed per mouse. For each mouse, 3 hypothalamic level-matched per mouse were used to quantify c-Fos immunoreactive cells in all AgRP and POMC immunostained cells observed in the ARC. The number of immunostained cells was counted manually using ImageJ software (Schneider et al., 2012) by an unbiased observer. For area measurements and particle counting, region of interest (ROI) within fluorescence images was manually selected with the mouse brain atlas for ARC, DMH and PVN, and was then measured by ImageJ software as previously described (Jin et al., 2016).

### Hypothalamic primary neuronal cell culture

Eight to ten neonatal (0-1 day old) pubs from either control and *Drp1*^*AgRPKO*^ mice derived from homozygous Cre-positive parents were used for hypothalamic primary neuronal cell culture. In brief, carefully removed the hypothalamus of the brain and placed onto a small culture dish that contains a small volume of Hibernate-A Medium (Cat# A1247501, Gibco). The tissues dissociated to single cells after digestion with 6 mL of Hibernate-A Medium containing 2.5% of Trypsin-EDTA for 15 minutes at 37°C. Suspended cells were filtered (40 µm) and centrifuged for 5 min at 1000 rpm and the pellet was re-suspended and plated on XF96 cell culture microplates (Cat# 101085-004, Agilent Technologies) coated with poly-D-lysine (Cat# P6407, Sigma) at a density of 1×10^5^ cells per well, and they were cultured in Neurobasal medium (Cat# 21103049, Gibco) supplemented with 1% penicillin-streptomycin, 2% B-27 Supplement (Cat# 17504044, Gibco) and GlutaMAX-I (Cat# 35050061, Gibco). After 10 days in culture, primary neuronal cells isolated from control and *Drp1*^*AgRPKO*^ mice were treated 2 µM 4-hydroxytamoxifen (H7904, Sigma-Aldrich) for expression of a CreER recombinase. Primary neuronal cells were used for the measurement of mitochondria fatty acid oxidation 5 Days later.

### Measurement of mitochondrial fatty acid oxidation assay

The fatty acid oxidation (FAO) was measured using a microfluorimetric Seahorse XF96 Analyzer (Agilent Technologies) according to the protocol supplied by the manufacturer with minor modifications. Cells were starved with minimal substrate neurobasal-A medium (Cat# 10888022, Thermo Fisher Scientific) for 24 hours. The minimal substrate medium included 1% B-27 Supplement (Cat# 17504044, Gibco), 1 mM glutamine, 0.5 mM carnitine, and 2.5 or 0.5 mM of glucose. The day of the assay, 45 minutes prior to the assay, starved cells were washed and incubated with Seahorse XF Base medium Minimal DMEM (Cat# 102353-100, Agilent Technologies) supplemented with 2.5 or 0.5 mM glucose and 0.5 mM carnitine in a non-CO_2_ 37°C incubator. 15 minutes prior to the assay, 40 µM etomoxir was added to the cells to measure endogenous fatty acid uptake for FAO. Palmitate-BSA or BSA control (Seahorse XF Palmitate-BSA FAO substrate, Cat# 1102720-100, Agilent Technologies) were added to cells right before initiating the XF assay. During the assay, cells were exposed to compounds in the following order: 5 µM of oligomycin (Cat# 495455, Sigma), 10 µM of FCCP [carbonyl cyanide-p-(trifluoromethoxy) phenylhydrazone] (Cat# C2920, Sigma), and 10 µM of antimycin A (Cat# A8674, Sigma) and 5 µM of rotenone (Cat# R8875, Sigma). Wave 2.6.0 (Agilent Technologies software, USA) software was used to analyze the parameters.

### Electrophysiology analysis

Electrophysiology analyses were performed as previously described (Toda et al., 2016). Briefly, eleven-to twelve-week-old mice were used for recordings. After mice were anesthetized with isoflurane and decapitated, the brains were rapidly removed and immersed in an oxygenated cutting solution at 4°C containing (in mM): sucrose 220, KCl 2.5, NaH2PO4 1.23, NaHCO3 26, CaCl2 1,MgCl2 6, and glucose 10, pH (7.3) with NaOH. After being amputated to a small tissue block, coronal slices containing the hypothalamus (300 µm thick) were cut with a vibratome. After preparation, slices were stored in a holding chamber with an oxygenated (with 5% CO2 and 95% O2) artificial cerebrospinal fluid (aCSF) containing (in mM): NaCl 124, KCl 3, CaCl2 2, MgCl2 2, NaH2PO4 1.23, NaHCO3 26, glucose 3, pH 7.4 with NaOH. The slices were eventually transferred to a recording chamber perfused continuously with aCSF at 33°C at a rate of 2 ml/min after at least a 1 h recovery in the storage chamber. Perforated patch recording was performed in AgRP-Tomato neurons of the ARC under voltage and current clamp. The membrane and spontaneous action potential were recorded in AgRP neurons under zero current clamp condition. For ghrelin-induced AgRP neuronal activation, baseline activity was recorded for at least 15 min. Slices were then perfused with 10 nM ghrelin, diluted in aCSF for 3 min, followed by a washout (with no ghrelin). At the end of the perforated patch recordings, the membrane of every cell was ruptured and whole-cell patch recording measured to check current-voltage relationship. All data were sampled at 5 kHz, filtered at 2.4 kHz, and analyzed with an Apple Macintosh computer using AxoGraph X (AxoGraph Scientific). Statistics and plotting were performed with KaleidaGraph (Synergy Software) and Igor Pro (WaveMetrics). The average firing rate was calculated in the last 2 min of each control period or treatment application. All the experiments were performed blindly to the electrophysiologist.

### Ghrelin administration

Individually housed 4 months old mice were i.p. injected with either 0.9 % saline (#0409-1966-12, Hospira Inc.) or ghrelin (10 nmol, HOR-297-B, Prospec) at 9:00 AM. Immediately after injection, mice were returned to their home cages, which contained a pre-weighed amount of food. The remaining food was measured at 0.5, 1, 2 and 4 hr post-injection. For immunostaining, mice were injected with ghrelin at 9:00 AM and 1 hr later, mice were deeply anesthetized and transcardially perfused, and brains were dissected and sectioned (50 µm) using a vibratome. Brain sections were processed for c-Fos immunostaining. Fluorescent images were captured with a Fluorescence Microscope (BZ-X710, KEYENCE). c-Fos/AgRP positive cells were counted using ImageJ software.

### Electron microscopy analysis

5-month-old Mice were deeply anesthetized and transcardially perfused with 0.9% saline containing heparin (10 mg/L), followed by fresh fixative (4% paraformaldehyde, 15% picric acid, 0.1% glutaraldehyde in 0.1 M PB). Brain coronal sections were immunostained with rabbit anti-RFP antibody (diluted 1:1,000 in 0.1 M PB, 600-401-379, Rockland) for AgRP neurons. After several washes with 0.1 M PB, sections were incubated with biotinylated goat anti-rabbit IgG (diluted 1:250 in 0.1 M PB, BA-1000, Vector Laboratories) for 2 hr at RT, then rinsed in 0.1 M PB three times 10 min each time, and incubated for 2 hr at RT with avidin–biotin–peroxidase (ABC; diluted 1:250 in 0.1 M PB; ABC Elite kit, Vector Laboratories). The immunoreaction was visualized with 3,3-diaminobenzidine (DAB). Sections were then osmicated (1% osmium tetroxide) for 30 min, dehydrated through increasing ethanol concentrations (using 1% uranyl acetate in the 70% ethanol for 30 min), and flat-embedded in araldite between liquid release-coated slides (Electron Microscopy Sciences). After capsule embedding, blocks were trimmed. Ribbons of serial ultrathin sections were collected on Formvar-coated single slot grids and examined using a Philips CM-10 electron microscope. Mitochondria morphology in AgRP neurons of fed and fasted mice were analyzed using ImageJ software as previously described (Toda et al., 2016).

### Measurement of circulating hormones

Five months old mice were deeply anesthetized and decapitated. The blood was collected into a capillary tube (Microvette, CB 300 Z, Sarstedt) containing 0.2 mg 4-(2-aminoethyl)-benzene-sulfonyl fluoride (AEBSF, Roche). Serum from blood samples was obtained by centrifugation at 3,000 rpm for 15 min, and each circulating hormones were determined using a commercially available ELISA kit for total ghrelin (Rat/Mouse Total Ghrelin ELISA kit, EZRGRT-91K, Millipore), and active ghrelin (Rat/Mouse Total Ghrelin ELISA kit, EZRGRT-90K, Millipore). All procedures were performed by following the manufacturer’s protocol.

### Statistical Analysis

Two-way ANOVA was used to determine the effect of the genotype and treatment with the Prism 7.01 software (GraphPad Software). For repeated measures analysis, ANOVA was used when values over different times were analyzed. When only two groups were analyzed, statistical significance was determined by an unpaired Student’s t-test. A value of p < 0.05 was considered statistically significant. All data is shown as mean ± SEM, unless otherwise stated.

## Acknowledgment

This work was supported by NIH R01 DK097566 to S.D.

