## Supplemental figures for "DRP1 IS REQUIRED FOR AGRP NEURONAL ACTIVITY AND FEEDING"

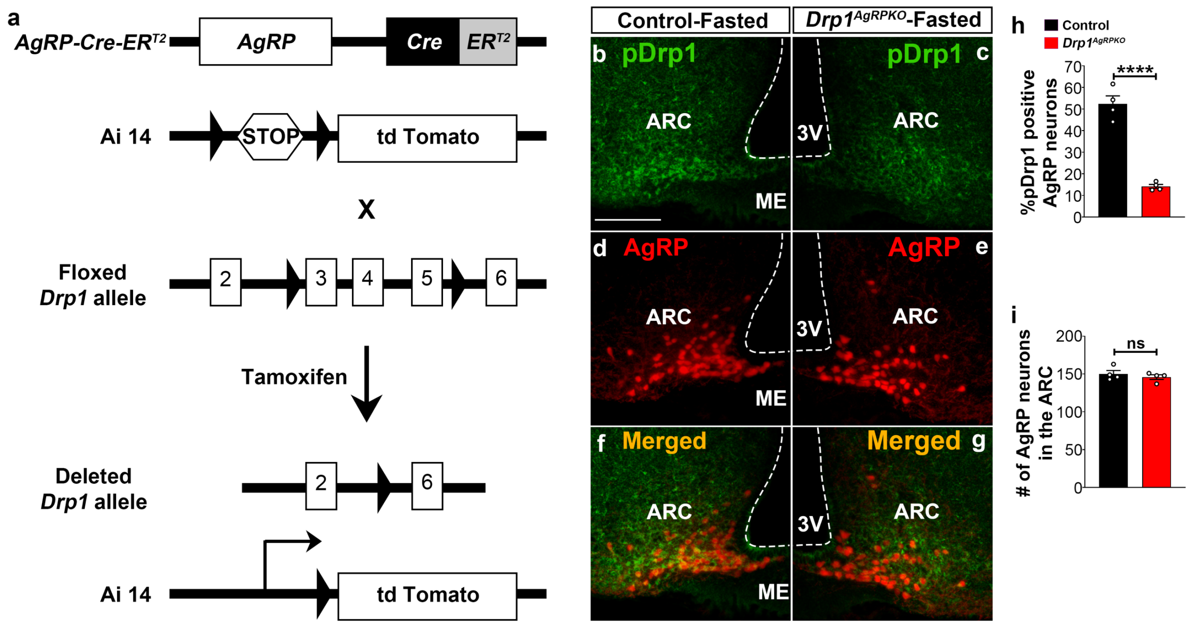


**Figure S1. Generation of AgRP neurons-specific *Drp1* deleted mice, related to Figures 3-7.**

**(a)** Schematic showing *Drp1*$^{fl/fl}$ and *AgRP-cre:ER*$^{T2}$ constructs and the strategy used to generate AgRP neurons-specific *Drp1* knockout mice. First, we generated *AgRP-cre:ER*$^{T2}$ mice harboring inducible *td*Tomato floxed by stop codon and then we crossed them with *Drp1*$^{fl/fl}$ mice.

**(b-g)** Representative micrographs showing immunostaining for phosphorylated Drp1 (at serine 616; pDrp1; green, b-c) and fluorescent reporter gene *td*Tomato (red, representing AgRP, d-e) and merged (f-g) in the hypothalamic ARC of a fasted control (b,d,f,) and a fasted *Drp1*$^{AgRPKO}$ male mouse (c,e,g). Scale bar represents 100 µm. 3v=third ventricle; ARC=arcuate nucleus; ME=median eminence.

**(h)** Graph showing quantification of pDrp1 expression in AgRP neurons of fasted control (n=4 mice) and *Drp1*$^{AgRPKO}$ male mice (n=4 mice). Data are presented as mean ± SEM. ****=P<0.0001 by unpaired two-tailed Student's t-tests.

**(i)** Graph showing no difference in total AgRP cell number between fasted control (n=4 mice) and *Drp1*$^{AgRPKO}$ mice (n=4 mice). Data are presented as mean ± SEM. P=0.4605 by two-tailed Student's *t*-test. ns=not significant.

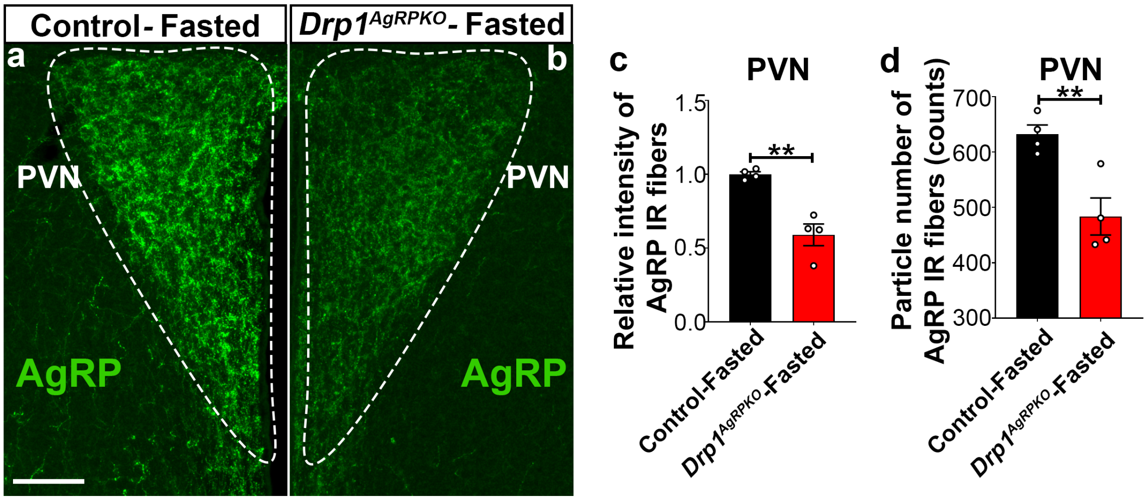


**Figure S2. AgRP-selective Drp1 deletion alters AgRP projection in female mice, related to Figures 4 and 5.**

**(a-b)** Representative micrographs showing immunostaining for AgRP (green) in the hypothalamic PVN of a fasted control (a) and a fasted $Drp1^{AgRPKO}$ female mouse (b). Dashed lines delineate the PVN. Scale bar represents 100 μm. PVN=paraventricular nucleus.

**(c-d)** Graphs showing quantification of relative intensity (c) and particle number (d) of AgRP fibers in PVN of fasted control and $Drp1^{AgRPKO}$ female mice (n=4 mice). Data are presented as mean ± SEM. **=P<0.01 by unpaired two-tailed Student's t-tests.

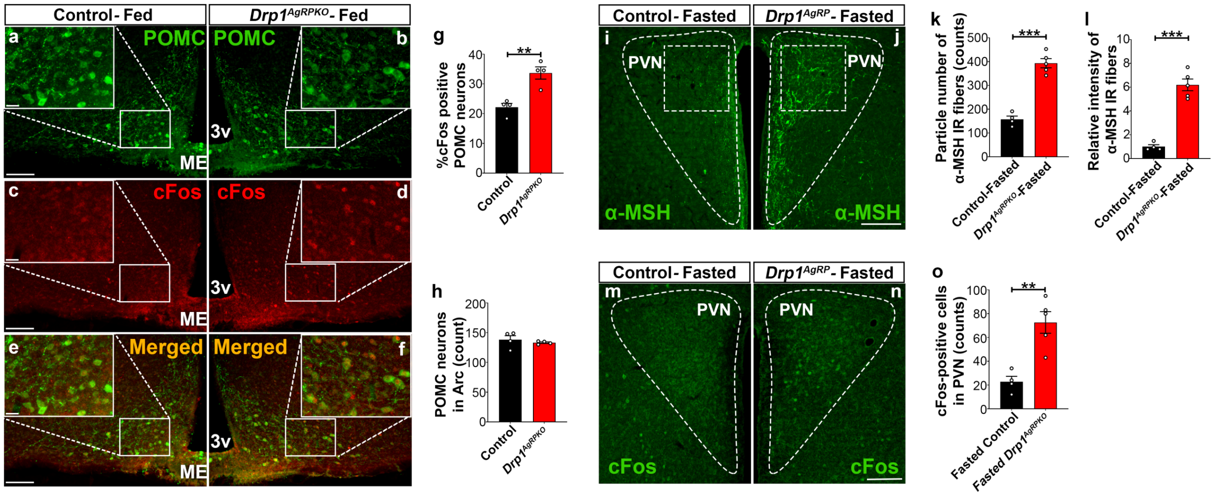


**Figure S3. Deletion of DRP1 in AgRP neurons affects hypothalamic POMC neurons, related to Figures 4 and 5.**

**(a-f)** Representative micrographs of hypothalamic sections showing immunostaining for POMC (green, a-b) c-Fos (red, c-d) and merged (e-f) in the hypothalamic ARC of a 5 months old fed control (a,c,e) and a 5 months old fed *Drp1$^{AgRPKO}$* male mouse (b,d,f).

**(g)** Graph showing the percent of c-Fos positive POMC neurons in 5 months old fed control (n=4 mice) and *Drp1$^{AgRPKO}$* male mice (n=4 mice). Data are presented as mean ± SEM. **=P<0.01 by two-tailed Student's *t*-test.

**(h)** Graph showing no difference in total POMC cell number between 5 months old fed control (n=4 mice) and *Drp1$^{AgRPKO}$* male mice (n=4 mice). Data are presented as mean ± SEM. P=0.4951 by two-tailed Student's *t*-test.

**(i-j)** Representative micrographs of hypothalamic sections showing immunostaining for α-MSH fibers in a fasted control (i) and a fasted *Drp1$^{AgRPKO}$* male mouse (j). Dashed lines delineate the PVN. Squared area represents the region used (ROI=region of interest) for analysis.

**(k-l)** Graphs showing quantification of relative intensity (k) and particle number (l) of α-MSH fibers in hypothalamic PVN of fasted control (n=4 mice) and *Drp1*$^{AgRPKO}$ male mice (n=5 mice).

**(m-n)** Representative micrographs of hypothalamic sections showing immunostaining for c-Fos from fasted control (m) and *Drp1*$^{AgRPKO}$ mouse (n).

**(o)** Graph showing quantification of c-Fos expression in hypothalamic PVN neurons of fasted control (n=4 mice) and *Drp1*$^{AgRPKO}$ male mice (n=5 mice). Data are presented as mean ± SEM. **=P<0.01; ***=P<0.001 by two-tailed Student's *t*-test. Scale bar represents 100 µm (a,d,g). Scale bar in high magnification image represents 20 µm (a). 3v= third ventricle; PVN=paraventricular hypothalamus; ME=median eminence.

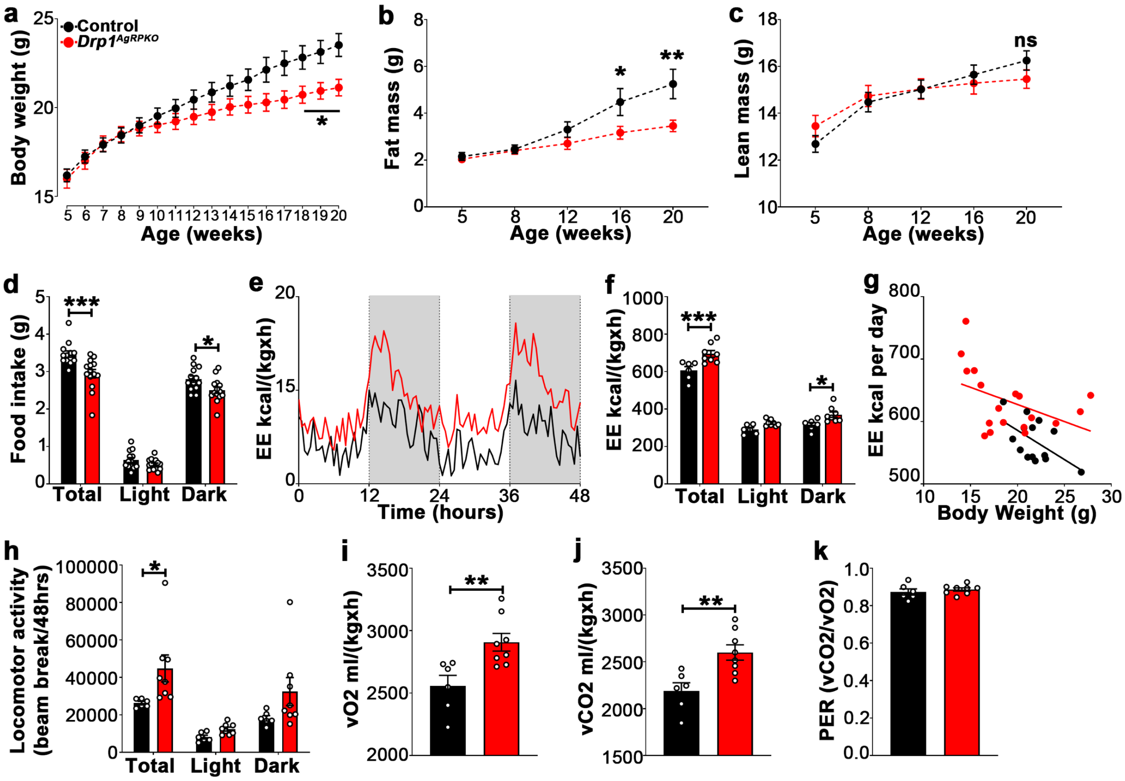


**Figure S4. Selective Drp1 deletion in AgRP neurons affects metabolic phenotype in female mice, related to Figure 6 and 7.**

(**a**) Graphs showing body weight of *Drp1*$^{+/+}$*-AgRP-cre:ER*$^{T2}$ mice injected with tamoxifen (n=20 mice) as control groups, and *Drp1*$^{fl/fl}$*-AgRP-cre:ER*$^{T2}$ mice injected with tamoxifen (n=22 mice). Data are presented as mean ± SEM. *P<0.05 by two-way ANOVA with Tukey's post hoc analysis for multiple comparisons.

**(b-c)** Graph showing fat mass (b), and lean mass (c) of *Drp1*$^{+/+}$*-AgRP-cre:ER*$^{T2}$ mice (n=11 mice) and *Drp1*$^{fl/fl}$*-AgRP-cre:ER*$^{T2}$ mice (n=10 mice). Data are presented as mean ± SEM. *P<0.05; **P<0.01 by two-way ANOVA with Tukey's post hoc analysis for multiple comparisons. ns=not significant.

**(d)** Graphs showing 24 hours food intake (average of 3 days) in 4 months old female control (n=13) and *Drp1*$^{AgRPKO}$ mice (n=14) and results of food intake as total in the 24 h cycle and in the dark and light phases of the cycle. Data are presented as mean ± SEM. *P<0.05; ***P<0.001 by two-way ANOVA with Tukey's post hoc analysis for multiple comparisons.

(**e-k)** Graphs showing energy expenditure (e-g), locomotor activity (h), consumed $O_2$ (i), produced $CO_2$ (j), and the respiratory exchange ratio (RER) (k) in 4 months old control and *Drp1*$^{AgRPKO}$ female mice. Gray area represents dark phases (e). Data are presented as mean ± SEM. *P<0.05; ***P<0.001 by two-way ANOVA with Tukey's post hoc analysis for multiple comparisons (f,h). P=0.9987 by linear regression analysis (g). **P<0.01 by two-tailed Student's *t*-test (i,j,k).

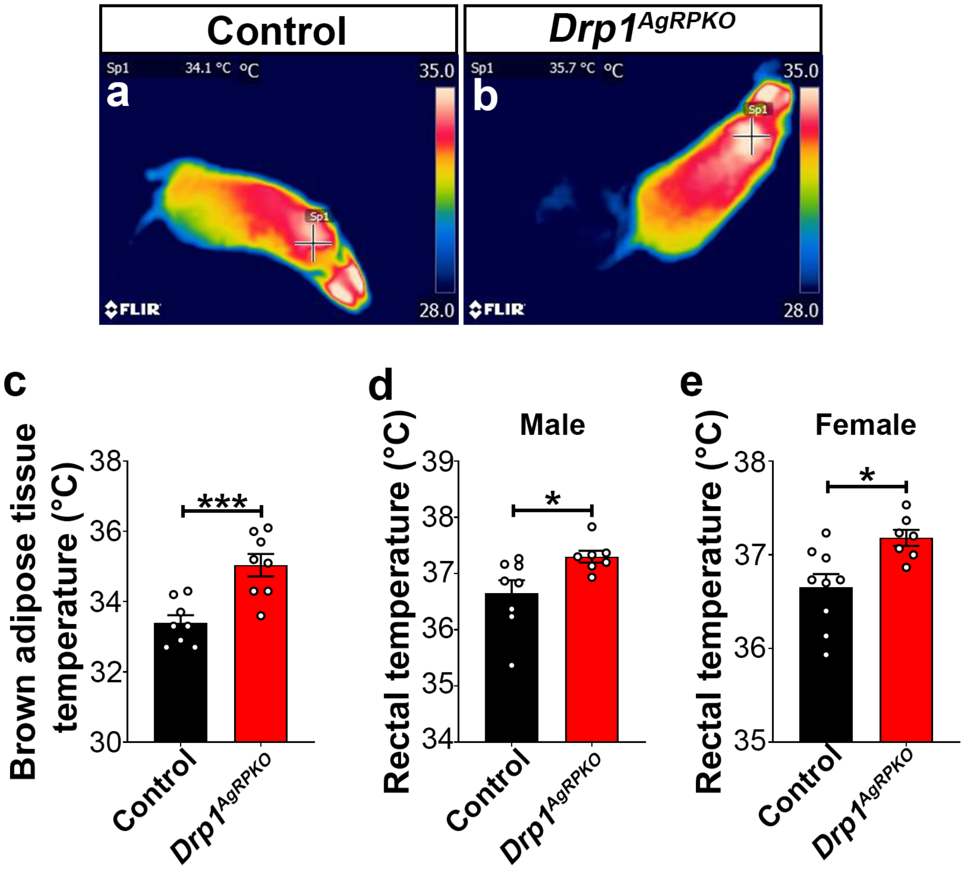


**Figure S5. Deletion of Drp1 in AgRP neurons results in increased BAT and core body temperature, related to Figures 6-7.**

**(a-b)** Representative infrared thermography images showing the temperature of the surface overlying BAT in control (a) and *Drp1*$^{AgRPKO}$ male mouse (b).

**(c)** Graph showing quantification of BAT temperature of the control (n=8 mice) and *Drp1*$^{AgRPKO}$ male mice (n=8 mice) at 4 months of age. Data are presented as mean ± SEM. ***=P<0.001 by unpaired two-tailed Student's t-tests.

**(d)** Graph showing rectal temperature in 4 months old control and *Drp1*$^{AgRPKO}$ male mice. Data are presented as mean ± SEM. *=P<0.05 by unpaired two-tailed Student's t-tests.

**(e)** Graph showing rectal temperature of 4 months old control and *Drp1*$^{AgRPKO}$ female mice. Data are presented as mean ± SEM. *=P<0.05 by unpaired two-tailed Student's t-tests.

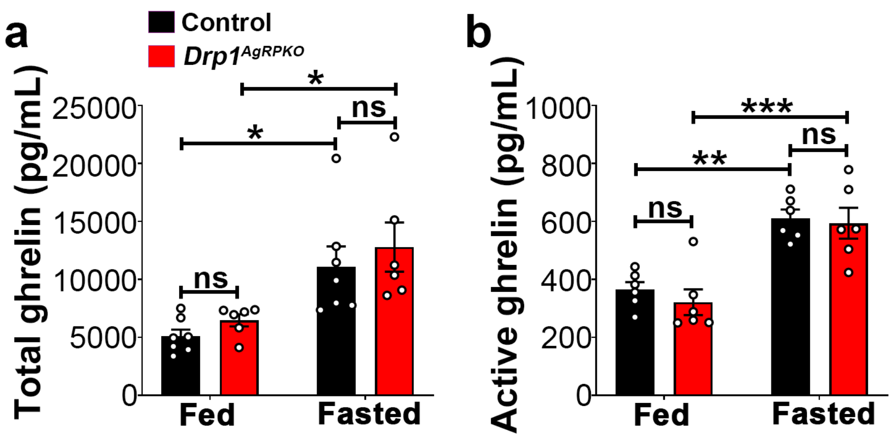


**Figure S6. Ghrelin levels in male mice, related to Figures 6-7.**

**(a)** Graph showing serum total ghrelin levels of 5 months old control male (n=7 mice) and *Drp1*$^{AgRPKO}$ male mice (n=6 mice) on fed and fasted states. Data are presented as mean ± SEM. *=P<0.05 by two-way ANOVA with Tukey's post hoc analysis for multiple comparisons. ns=not significant.

**(b)** Graph showing serum active ghrelin levels of 5 months old control male (n=6), and *Drp1*$^{AgRPKO}$ male mice (n=6) on fed and fasted states. Data are presented as mean ± SEM. **=P<0.01; ***=P<0.001 by two-way ANOVA with Tukey's post hoc analysis for multiple comparisons. ns=not significant.
